# Nanoscale insights into chromatin integrity molecular rearrangements upon DNA damage response

**DOI:** 10.64898/2026.09.13.751252

**Authors:** Sara Seweryn, Karolina Juzba, Michał Czaja, Katarzyna Skirlińska-Nosek, Marta Urbańska, Anna Nowakowska, Wojciech Kwiatek, Michał Sarna, Marek Szymoński, Bayden Wood, Ewelina Lipiec

## Abstract

DNA Double Strand Breaks (DSBs) threaten genomic stability, leading to cell death, chromosomal rearrangements, and cancer-driving mutations. Therefore, the effective repair mechanisms are essential for maintaining genomic stability and ensuring cellular survival across diverse organisms. At the core of this process lies chromatin integrity, which facilitates the local DNA conformational changes, regulating accessibility to repair proteins and other biomolecules. To deepen our understanding of the DNA damage response pathway, the local molecular mechanisms regulating the interplay between DNA damage formation and alterations in chromatin conformation must be investigated at the nanoscale level. Here, we integrated atomic force microscope-infrared spectroscopy (AFM-IR) and confocal fluorescence microscopy to explore local chemical modifications in DNA structure and chromatin integrity in metaphase chromosomes and chromosomal aberrations isolated from cells treated with the chemotherapeutic agent, bleomycin. Our findings reveal changes in secondary protein structures, indicating the engagement of DNA repair proteins with high β-sheet content. Nanospectroscopic mapping resolved the alterations in DNA condensation along the chromosomes. Moreover, we observed global DNA demethylation, particularly the conversion of 5-methylcytosine (5mC) to 5-hydroxymethylcytosine (5hmC), correlating with increased DSBs. We conclude that these transitions in protein conformation and DNA methylation correlate with chromatin relaxation and enhanced accessibility for the repair protein.

**GRAPHICAL ABSTRACT:** 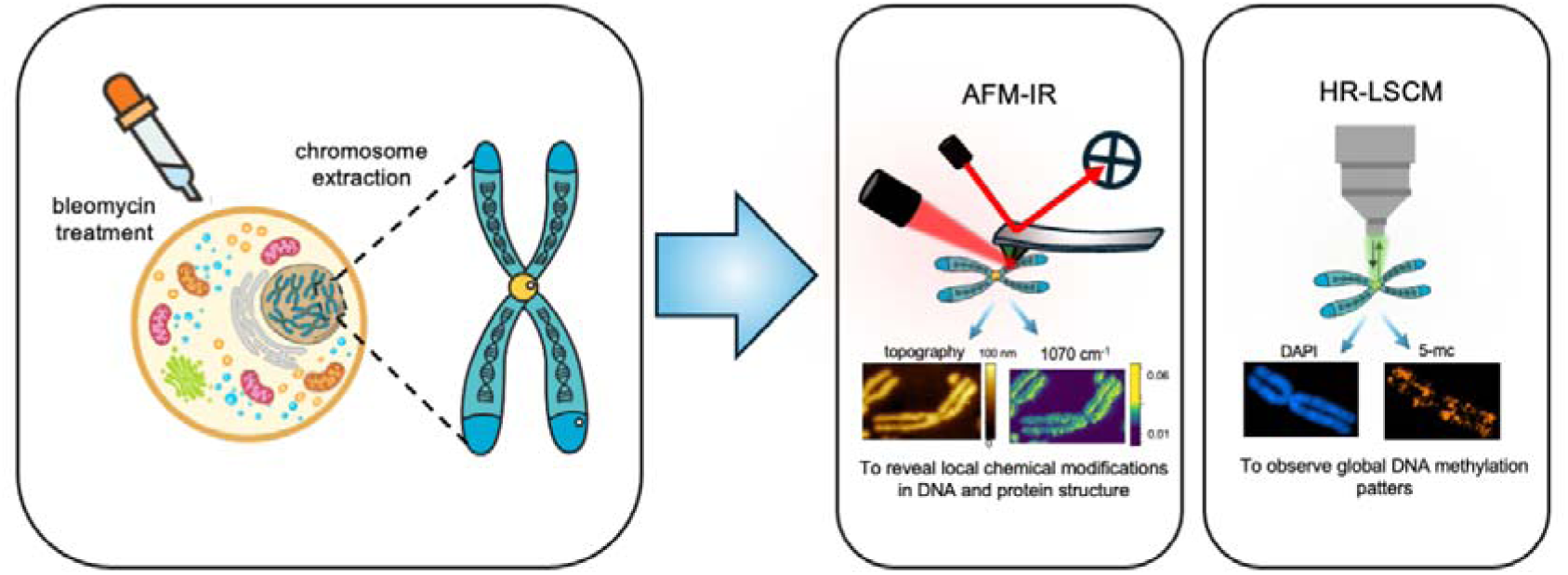

## INTRODUCTION

Cellular DNA is constantly exposed to many damaging factors, these include exogenous agents such as ionising radiation or chemotherapeutic drugs, and free radicals generated by endogenous cellular processes [1]. Among many types of DNA damage, Double Strand Breaks (DSBs) are the most harmful DNA lesions. DSBs sever phosphate backbones of the complementary DNA strands and compromise genome integrity, disrupting genetic information, nuclear architecture, and local chromatin structure. An inability to properly respond or repair, DSBs trigger the response that leads to cell death and oncogenic mutations [2]. Therefore, cells have evolved multiple complex mechanisms to repair DNA damage, essential for preserving genomic integrity across all life forms.

The effectiveness of numerous cancer therapies relies on inducing DSBs to eliminate uncontrolled and rapidly proliferating tumour cells. One of the well-known exogenous radiomimetic agents is the chemotherapeutic drug Bleomycin (BLM), clinically used in anticancer treatments for various types of cancer (i.e., testicular germ-cell tumour, squamous cell carcinoma) [3], [4]. In the presence of metal ions (Cu(II) and Fe(II)) and oxygen, BLM binds to DNA and induces both single and double strand breaks, with the double strand breaks being considered the primary cause of cytotoxicity [5]. In mammalian cells, approximately 70-85% of DSBs are repaired through the non-homologous end joining (NHEJ) pathway, which constitutes the dominant repair mechanism throughout most phases of the cell cycle In the context of bleomycin-induced damage, several studies have demonstrated that DSB repair is predominantly mediated by NHEJ [6], [7]. Mladenov et al. showed that in human HeLa cells treated with BLM, homologous recombination (HR) markers such as RAD51 foci appear at later stages of the damage response, particularly when cells accumulate in the G2 phase of the cell cycle, indicating a secondary role of HR when NHEJ is saturated or inefficient [8]. Consistently, cellular sensitivity studies have revealed that deficiency in key NHEJ components (e.g, DNA-PKcs, LIG4, 53BP1) increases susceptibility to bleomycin-induced cytotoxicity, further supporting the predominant role of NHEJ in repairing BLM-induced DSBs [9]. It has been determined that BLM preferentially targets 5′-GT and 5′-GC dinucleotides. The cleavage of DNA strands starts with the interaction of BLM with DNA (either through minor groove binding or intercalation), which is then followed by the targeting of pyrimidine nucleotides (T, C) [10], [11]. Numerous in vitro experiments performed on DNA solutions and live cells have verified the efficacy of this glycopeptide antibiotic, although its impact on local DNA conformational transition, followed by the DNA repair process, is still poorly understood.

It is becoming increasingly evident that DNA conformational transitions are implicated in both DNA susceptibility and repair processes [12]. The structural stability of the DNA double helix relies on hydrogen bonds formed between nitrogenous bases on the opposite strands, enabling the molecule to adopt various secondary structures. The majority of DNA in functional cells exists in the B-DNA form, with the capacity to also fold into A-DNA and Z-DNA conformations, each differing in the dimensions of their major and minor grooves [13], [14]. A-DNA has more base pairs (bp) per helical turn compared to B-DNA, leading to increased compression along its axis. Similarly, a reduction in length by approximately 20–25% is observed for A-DNA compared to B-DNA with an equivalent number of base pairs [13]. Sequences characterised by alternating purine/ pyrimidine bases promote the left-handed Z-DNA conformation [15]. The structural properties of DNA are crucial in determining its accessibility to interact with other macromolecules. The presence of locally induced A-DNA conformation has already been observed in some protein-DNA complexes [16]. Additionally, it is postulated that the local conformational transition from B-DNA to A-DNA occurs within regions of DNA damage [13], [17].

In eukaryotic cells, the repair of DSBs occurs within a highly dynamic and complex chromatin environment [18]. Inside the nucleus, the chromatin structure segregates spatially into heterochromatin and euchromatin regions. Euchromatin refers, in general, to genome regions containing actively transcribed genes or potentially active ones, which undergo decondensation during interphase [19]. Heterochromatin is a highly condensed DNA structure, containing approximately 20% of the mapped human genome. It is characterised as transcriptionally inactive but essential for maintaining structural integrity [20], [21]. The main chemical difference between these two forms of DNA is the density of pyrimidines and purines methylation, which increases from the euchromatin towards the heterochromatin. Chemically, DNA methylation involves the substitution of a methyl group to the C5 position of the cytosine (5mC), occurring in regions preceding guanine nucleotides or CpG dinucleotides (cytosine (C) and guanine (G) are separated by a single phosphate group (p)). Usually, linear CpG sequences are methylated on both strands of DNA [22], [23]. The CpG-rich regions are frequently located at gene promoters, rendering them susceptible to aberrant methylation. For instance, elevated CpG methylation level results in a higher incidence of spontaneous DNA mutations, as the deamination of methylated cytosines (mCs) converts them into thymine, creating a T:G mismatch [24]. Another DNA modification is cytosine hydroxymethylation (5hmC), proposed as a new epigenetic marker of DNA damage [25]. 5hmC is a DNA base formed by active oxidation of 5mC by TET dioxygenases during the DNA demethylation process [26]. Subsequent studies have demonstrated that 5hmC acts as a stable epigenetic regulator of gene expression and that DNA containing 5hmC typically presents an open chromatin structure [27], [28]. Research has shown that 5hmC accumulates at sites of DNA damage, induced by aphidicolin and microirradiation, colocalising with DNA repair proteins 53BP1 and γH2AX [29]. DNA methylation has an essential role in regulating gene expression, cellular functionality, and genome integrity. Consequently, it affects not only the gene activity but also the mechanical properties of DNA [30], [31], [32], which influence chromatin activities such as the interactions between DNA and proteins [33], [34], and other molecules such as drugs and dyes, and the resistance to strand separation [35].

In this study, we applied a combination of atomic force microscope-infrared spectroscopy (AFM-IR) and high-resolution laser scanning confocal microscopy (HR-LSCM) to explore chemical modifications of DNA structure and chromatin integrity in mechanisms of DNA damage and repair. In our previous work, we demonstrated that nanoscale vibrational spectroscopy enables direct tracking of DNA methylation patterns within native chromatin, allowing the spatial mapping of euchromatin and heterochromatin based only on their chemical signatures without any chemical labelling [36].To further extend this approach, we isolated individual metaphase chromosomes from cells treated with bleomycin, an anticancer agent known to induce DNA double-strand breaks [37], [38]. Metaphase chromosomes provide a highly condensed and structurally well-defined chromatin state, in which regions of higher and lower DNA methylation are spatially segregated, making them particularly suitable for studies of epigenetic modifications [38], [39]. All the investigated cells were in the metaphase stage of the cell cycle upon the ongoing repair processes. In case of a lethal number of DNA damage, the cell cycle stops at the G0 phase, and the development of the metaphase is largely prohibited [40]. To comprehensively analyse the chemical composition of metaphase chromosomes, we adopted a systematic approach, encompassing AFM mapping of the spatial distribution of spectral markers related to DNA damage and repair, including methylene, amide, and phosphate vibrations, as well as single-spectrum acquisition along a single chromosome coupled with multivariate data analysis. Furthermore, we performed fluorescence microscopy imaging of 5mC and 5hmC in chromosomes to identify chromatin methylated upon DNA damage formation through interaction with bleomycin.

## MATERIAL AND METHODS

### Bleomycin preparation

Bleomycin sulphate was procured from TCI Europe N.V. (Tokyo Chemical Industry). To prepare activated bleomycin for all experiments, ferric ammonium sulphate dodecahydrate was first dissolved in water, followed by the addition of a 10% molar excess of iron to bleomycin [41]. The 1 mM bleomycin-iron solution was then maintained at −20 °C for future use.

### Cell culture

Human cervical cancer cells (HeLa cell line), were cultured in a humidified incubator at 37°C in 5% CO2, in high glucose Dulbecco’s Modified Eagle Medium (DMEM, Gibco, glucose concentration of 4500 mg/L) supplemented with 10% Fetal Bovine Serum (FBS, Gibco) and 1% Penicillin/Streptomycin solution (Gibco).

Human lymphocytes were obtained from peripheral blood collected from a healthy, anonymous male donor (<35 years old) into heparin containing tubes to prevent coagulation. All procedures involving human material were conducted in accordance with relevant ethical guidelines and were approved by the local Bioethics Committee (approval no. 124/KBL/OIL/2013 and letter no. OIL/KBL/23/2018).

### Bleomycin treatment

HeLa cells were incubated with bleomycin, which was added directly to the cell culture medium at final concentrations of 50 and 150 μM. Cells were exposed to the drug for 24 and 48 hours to induce DNA DSBs. Following the incubation period, cells were processed for metaphase chromosome isolation in order to investigate the cellular response to drug-induced DNA damage. The applied concentrations and incubation times were selected based on previous studies, indicating that the 50 and 150 μM BLM concentrations prompted cellular repair mechanisms [42], [43].

### Ionising radiation exposure

Freshly collected human blood samples containing lymphocytes were transferred to 2 mL Eppendorf tubes immediately prior to irradiation and kept on ice before and after exposure. Samples were irradiated with 60 MeV proton beam at the Cyclotron Centre Bronowice (Institute of Nuclear Physics, Polish Academy of Science, Krakow, Poland). Irradiations were performed using a spread out Bragg peak (SOBP) configuration, with samples positioned at the isocenter of the irradiation setup inside a dedicated PMMA phantone equipped with an additional PMMA plate corresponding to a water-equivalent thickness of 4.72 mm H_2_O [44]. Blood samples were irradiated with doses of 3Gy. After 46h post irradiation, lymphocytes were processed for metaphase chromosome preparation.

### Chromosome preparation

Metaphase chromosome spreads were prepared from HeLa cells and human lymphocytes according to the standard cytogenetic procedure [45]. Briefly, after completion of drug treatment, cells were treated with colcemid (5%, Sigma Aldrich for HeLa cells; GIBCO for lymphocytes) to increase the population of actively dividing cells arrested in the metaphase stage. HeLa cells were incubated with colcemid for 4h, whereas lymphocytes were incubated with colcemid for 2h. Subsequently, cells were harvested (by trypsinisation in the case of adherent HeLa cells or by centrifugation for suspension lymphocytes) and subjected to hypotonic treatment for 10 mins to induce cell swelling ( 1 mM NaCl for HeLa cells or 0.075M KCL for lymphocytes), followed by centrifugation. The swollen cells were then fixed using a mixture of freshly prepared methanol (Sigma Aldrich) and acetic acid (Sigma Aldrich) in a 3:1 ratio. Aliquots of the fixed cell suspensions were deposited onto gold-coated silicon wafers (HeLa cells) or ZnS substrates (lymphocytes) for AFM-IR measurements (20 µl) and onto glass coverslips for immunofluorescence staining (20-40 µl). Upon contact with the substrate, the swollen cells ruptured, releasing metaphase chromosomes suitable for further analysis.

### Cellular nuclei isolation

Cellular nuclei were isolated following the protocol by Junaid et al. [46]. Cells were detached from culture flasks through trypsinisation and then treated with a nuclear extraction buffer containing 320 mM sucrose (84097, Sigma-Aldrich, purity >99.5% HPLC), 5 mM MgCl2 (84097, Sigma-Aldrich), 10 mM HEPES (H3375, Sigma-Aldrich, purity >99.5% titration), and 1% Triton X-100. The samples were then gently mixed using a vortex, incubated on ice for 10 minutes, and subsequently fixed with 4% formaldehyde. Before conducting SR-FTIR measurements, the suspensions were centrifuged to eliminate the buffer, and then nuclei were resuspended in physiological saline solution. Around 40 μl of each suspension was placed between two CaF_2_ slides and mounted in a custom-made sample holder [47].

### AFM IR spectra collection and AFM IR mapping

AFM-IR spectra and hyperspectral maps were acquired using a commercially available nanoIR2 microscope (BrukerTM, Anasys, Santa Barbara, CA, USA). Measurements were performed in contact mode using gold-coated silicon probes PR-EX-nIR2-10 (20 nm tip apex diameter, Res. Freq. 13±4 kHz, Spring. const. 0.07-0.4 N/m), using an Anasys instrument (BrukerTM). In all experiments, AFM topography images of metaphase chromosomes were collected before infrared measurements to identify regions of interest.

AFM-IR measurements on metaphase chromosome spreads prepared from HeLa cells were performed using a tunable infrared Quantum Cascade Laser (QCL). Following topographical imaging, chromosome I (representing ±8% of the total DNA in cells) was identified and selected for AFM-IR analysis. Spatial distribution of the absorption at 1070 cm^−1^, 1408 cm^−1^, 1540 cm^−1^ and 1660 cm^−1^ corresponding to peaks from phosphate motions of the DNA backbone, methylene groups, amide II and amide I vibrations, respectively, were mapped with a pixel size of 15 nm. All AFM-IR spectra were collected within the spectral range of 1712 - 1350 cm^-1^ and 1230-900 cm^-1^ with 1024 scans co-added. The scan areas corresponded to chromosome sizes, typically several microns, with scan rates adjusted between 0.02 Hz and 0.01 Hz. The spectral resolution was set to ∼2 cm^-1^ over the full laser spectral range.

AFM-IR measurements on metaphase chromosome spreads prepared from human lymphocytes were performed using the same nanoIR2 system operated with a tunable Optical Parametric Oscillator (OPO) laser in resonant contact mode. IR maps were acquired over areas of 2-10 μm, depending on chromosome size, with a spatial resolution of 250 – 450 pixels and scan rates between 0.01 and 0.02 Hz. AFM-IR maps were collected at 2960 cm^-1^ (methyl group), 1660 cm^-1^ (amide I), and 1240 cm^-1^ (asymmetric stretching of phosphate motions from the DNA backbone). AFM-IR spectra were recorded in the spectral ranges of 3100-2800 cm^-1^ and 1800-900 cm^-1^ with 1024 scans co-added and a spectral resolution of 4 cm^-1^ across the full laser tuning range.

AFM-IR measurements on metaphase chromosomes isolated from the MDA-T68 thyroid cancer cell line were performed using a nanoIR2 system equipped with a tunable infrared quantum cascade laser (QCL). AFM-IR maps were acquired at a spatial resolution of 300–500 pixels and scan rates of 0.2–0.3 Hz. The mapped region ranged from 3 to 8 μm, depending on the size of chromosome I. AFM-IR maps were collected at 2960 cm ¹ (methyl groups), 1660 cm ¹ (amide I), and 1240 cm ¹ (asymmetric stretching of phosphate groups in the DNA backbone). AFM-IR spectra were recorded over the spectral ranges of 3100–2800 cm ¹ and 1800–900 cm ¹, with 256 co-added scans and a spectral resolution of 4 cm ¹.

### Synchrotron FTIR (SR-FTIR) measurements

Infrared micro-spectroscopy experiments were conducted at the SISSI-Bio Beamline at Elettra Sincrotrone Trieste [48]. The measurements were performed using a Vertex 70v FTIR spectrometer coupled with a Hyperion 3000 IR microscope (Bruker), featuring a 15× Schwarzschild IR objective and a liquid nitrogen-cooled mercury cadmium telluride (MCT) detector. Spectra were acquired in transmission mode, averaging 512 scans at a scanner speed of 120 kHz with a spectral resolution of 4 cm ¹, covering the spectral range of 5000 to 600 cm ¹. A buffer reference spectrum was collected after every 10 spectra. Due to the high collimation and intensity of the synchrotron IR source, spectra were collected from a diffraction-limited spot with the square aperture set to 18 μm for nuclei.

### Spectral data processing for AFM-IR data

Single point spectra were corrected for the baseline (with 3rd order of polynomial), smoothed using Savitzky-Golay algorithm (2nd order of polynomial, 13 smoothing points) and normalised (Standard Normal Variate) using MATLAB R2019b software (MathWorks, Inc., USA). An average AFM-IR spectrum was then calculated for each treatment condition. Analysis and processing of AFM and infrared maps were conducted with SPIP software (Image Metrology, Denmark). Obtained images were normalised to the applied laser power, and their positions were correlated based on topographies that were collected simultaneously with each absorption map. The z-offset for each map was adjusted to set the minimum value to 1. For metaphase chromosome spreads prepared from HeLa cells, ratio maps were calculated by dividing the IR absorption map at 1408 cm^-1^ by the IR map at 1070 cm^-1^, and the IR absorption map at 1660 cm^-1^ by the IR map at 1540 cm^-1^. For chromosome spreads prepared from human lymphocytes exposed to ionising radiation, ratio maps were calculated by dividing the IR absorption map at 2960 cm^-1^ by the IR map at 1240 cm^-1^.

### Spectral data processing for SR-FTIR data

The SR-FTIR spectra were processed using signal smoothing (Savitzky-Golay algorithm, 3rd order of polynomial, 7 smoothing points) and baseline correction (2nd or 3rd order polynomial) through the built-in tools of OPUS software. To eliminate water interference, the buffer spectrum was multiplied by a contribution factor (ranging from 0.8 to 0.9) and subtracted individually from each analysed cell spectrum. An average FTIR spectrum was then calculated from 3 randomly selected spectra for each treatment condition. To better represent spectral variations in the DNA phosphate motions, the second derivative of the mean spectra in the 1200-1000 cm ¹ range was calculated using Origin Pro 2023 software.

### Multivariate Data Analysis

Principal Component Analysis (PCA) and k-means clustering were performed using MATLAB R2019b software (MathWorks, Inc., USA) and its built-in toolboxes.

PCA was conducted on the single-point AFM-IR spectra to reduce the high dimensionality of the spectral dataset by identifying key patterns and summarising the variance across the spectra into a smaller number of components. In this case, two principal components (PCs) were retained, capturing the key spectral features and simplifying the data while preserving most of the variance, thus facilitating the interpretation of differences and revealing underlying spectral variations not visible in the raw data.

The k-means clustering was applied to the hyperspectral maps to classify the data into distinct groups based on spectral similarity. The k-means algorithm was set to produce three clusters, where each cluster represented a unique combination of spectral features. The number of clusters (k = 3) was chosen based on preliminary analysis, reflecting the presence of three main spectral patterns in the data. Each pixel in the hyperspectral maps was assigned to one of these three clusters, allowing the generation of false-colour maps. These maps visually represented the spatial distribution of the clusters, with different colours corresponding to the different spectral groupings. This clustering approach helped to reveal spatial patterns and variations within the sample, offering insights into the heterogeneity of the material being studied.

### Immunofluorescent staining and imaging

The cell suspension was dropped onto glass coverslips and subjected to a staining protocol starting with chromosome denaturation in 4M HCl solution for 8 minutes. After denaturation, the samples were carefully rinsed with 0.1% Triton X (Sigma-Aldrich) dissolved in PBS at 43 ℃. Subsequently, chromosomes were incubated with blocking solution (1% BSA, 0.1% Triton X in PBS) for 30 minutes. A mixture of primary antibodies against 5hmc (5-Hydroxymethylcytosine Recombinant Rabbit Monoclonal Antibody RM236, Thermo Fisher Scientific, MA5-24695) and 5mc (5-Methylcytosine Monoclonal Antibody 33D3, Thermo Fisher Scientific, MA5-38432), diluted in blocking solution at a ratio of 1:250, was put on the slides in a humidity chamber for 90 minutes at room temperature. Then, the slides were rinsed again with 0.1% Triton X dissolved in PBS at 43 ℃ and a secondary antibody mixture (anti-mouse and anti-rabbit) diluted in blocking solution at a ratio of 1:250 was applied to the samples in a humidity chamber and incubated at 4 ℃ for 90 minutes. After incubation with the secondary antibodies, the slides were rinsed again with 0.1% Triton X dissolved in PBS at 43℃. Chromatin was then stained with Hoescht (Sigma-Aldrich) staining solution for 15 minutes, followed by a final rinse with 0.1% Triton X dissolved in PBS at 43 ℃. Excess of liquids was removed carefully without the samples being shrivel up. The slides were sealed by using Fluorescence Mounting Medium (Agilent Technologies, Inc., USA) with cover glass and imaged.

Fluorescence images were acquired using high resolution laser scanning confocal microscope (HR-SLCM) LSM 900 with Airyscan 2 from Zeiss. The microscope was equipped with four lasers: 405 nm, 488 nm, 561 nm and 640 nm, two multialkali detectors and one 32-channel GaAsP array detector. Images were obtained using a Plan-Apochromat 63x oil immersion objective with a numerical aperture of 1.4. Image analysis and processing were carried out using ImageJ software (1.54f, National Institutes of Health, USA).

DNA damage, visualised as fluorescent green 5-hydroxymethylcytosine (5hmC) puncta in chromosome images, was quantified using a custom ImageJ macro with manual adjustments for precision. Initially, chromosome 1 was delineated from each chromosome set in the images through freehand selection. The background was then subtracted, and a mean filter with a radius of 1 pixel was applied to enhance the visualisation of discrete puncta. Next, automated thresholding was applied, and the image was converted into a binary mask. The watershed function was used to separate connected puncta, and remaining unresolved clusters were manually segregated to ensure accurate quantification. Individual puncta, representing sites of DNA damage, were then counted using the “Analyse Particles” tool in ImageJ Fiji.

In parallel, comparative analysis of 5-methylcytosine (5mC) fluorescence was conducted to assess methylation patterns. For each selected chromosome I, the area occupied by the orange 5mC signal was measured using ImageJ. A fixed threshold value, determined through prior analysis of control images, was applied to consistently segment the 5mC signal, and the area was quantified using the “Measure” function. The number of counts and Area of 5-methylcytosine [a.u] values were statistically analysed by Two-way ANOVA and HSD Tukey test (P < 0.001). This analysis was carried out in the statistical program Statistica 13, 1984-2017 TIBCO Software Inc.

### Multicolour FISH (mFISH) chromosome identification

Multicolour fluorescence in situ hybridisation (mFISH) was performed on metaphase chromosome spreads prepared from human lymphocytes using a commercially available 24XCyte Human Mulicolor FISH probe set (MetaSystems, D-0125-060-DI). This probe set enables simultaneous identification of all 24 human chromosomes using chromosome-specific fluorescent DNA probes [49]. Briefly, chromosomal DNA was denatured, followed by hybridisation with the mFISH probe mixture under controlled temperature and humidity conditions according to the manufacturer’s protocol. After hybridisation, a series of post-hybridisation washes was performed to remove unbound probes, and chromosomes were counterstained with DAPI. Fluorescence images were acquired using a MetaSystem imaging system and ISIS software.

To ensure statistical robustness and reproducibility, the analysis was conducted across a large number of metaphases, and the data were compared across independent biological replicates (at least three). Figures presenting individual metaphases are representative, whereas semiquantitative analyses of DNA methylation and multivariate statistical analyses are based on data acquired from multiple biological replicates.

## RESULTS AND DISCUSSION

### AFM-IR spectra of individual metaphase chromosomes and FTIR microspectroscopy of isolated cellular nuclei exposed to an anticancer drug

AFM-IR spectra of chromosome I were compared with spectra of cellular nuclei obtained using SR-FTIR (Synchrotron Radiation Fourier Transform Infrared Spectroscopy) acquired at the Elettra Sincrotrone Trieste. This allowed a direct comparison of spectral changes related to BLM treatment at the chromosome and cellular nuclei level. SR-FTIR spectra acquisition from individual chromosomes is prohibited due to their small size. Signal-to-noise ratio and lateral spatial resolution of ∼2.5 µm achievable with conventional infrared spectroscopy methods [50] are not sufficient enough for the acquisition of spectra from objects as small as individual chromosomes (thickness 100-200 nm). Additionally, the sample preparation protocol precludes the preparation of a bulk solution containing exclusively chromosomes, as they are released from nuclei only upon contact with the substrate surface [45]. Figure 1 presents a comparison of AFM-IR spectra (60 averaged spectra acquired from two different chromosomes) of chromosomes isolated from the control group and bleomycin-treated cells, with spectra of cellular nuclei (3 averaged spectra) collected using a synchrotron light source. Before the cellular nuclei isolation, cells were treated with the same experimental conditions as used for chromosome measurements. As expected, the chromosome spectra closely resemble the cellular nuclei spectra. Typical spectral features are observed, including symmetric stretching of phosphate from the DNA backbone in the spectral range 1090 - 1060 cm^-1^ [51], amide I and amide II vibrations in the spectral ranges: 1690 - 1600 cm^-1^ and 1590 - 1500 cm-1, respectively [52], [53] and methyl and methylene bending motions at 1450 cm^-1^ [52]. The spectral shifts observed in AFM-IR vibrational modes, in comparison to standard ST-FTIR spectra, are due to the influence of wavelength-dependent reflectivity and thermal response of the substrate underneath [54], [55]. It is worth noting that the SR-FTIR spectra exhibit baseline variations in comparison to the AFM-IR data. This discrepancy may be attributed to variability in optical properties of individual cells, including thickness, the distribution of the local density and possibly the Mie scattering, as the size of the nuclei is similar to the wavelengths of IR radiation, particularly affecting the 1200-1000 cm ¹ spectral region [56]. In AFM-IR, thermal expansion in the sample is detected therefore spectral shape (and baseline) is not affected by any type of scattering. Following a cellular response to DNA damage, characteristic changes associated with the protein expression and conformational modifications of DNA resulting from bleomycin treatment are observed.

**Figure 1.**
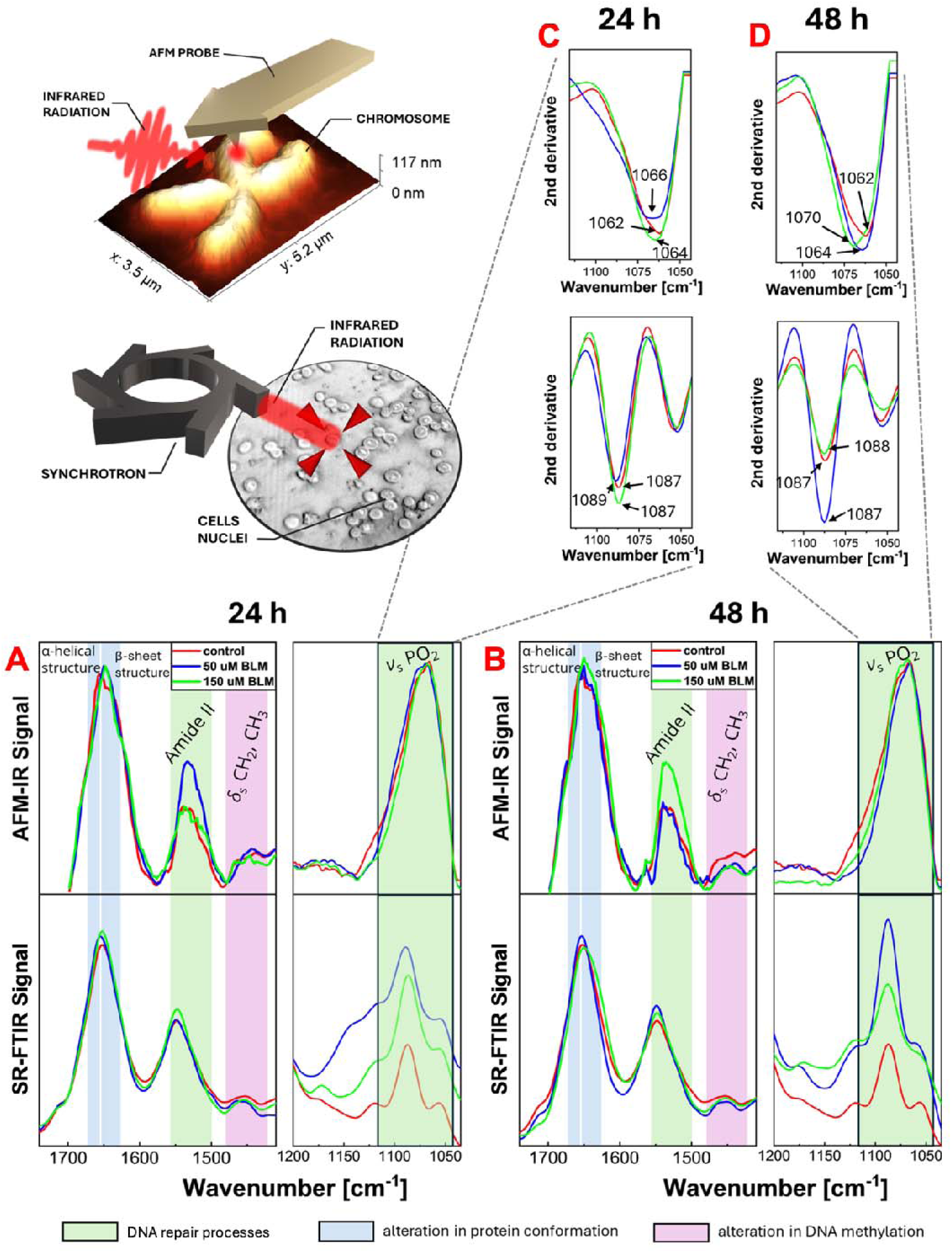
Averaged infrared spectra illustrating DNA chemical modifications induced by bleomycin treatment. (A,B) The averaged AFM-IR spectra of single metaphase chromosomes I were compared with the averaged SR-FTIR spectra of isolated cellular nuclei after 24h and 48h bleomycin treatment. (C,D) Second derivative spectra corresponding to panels A and B, calculated to minimise baseline effects and enhance subtle spectral differences. The shaded region highlights the 1100-1050 cm^-1^ spectral range associated with phosphate vibrations of the DNA backbone.

Spectral changes, such as intensity variations within the amide I and II regions and shifts in the spectral position associated with phosphate motions from the DNA backbone νs (PO2^-^) are characteristic markers of DNA repair processes. Specifically, an increase in amide II intensity indicates a cellular response to DNA damage, leading to an increased expression of repair proteins [57]. Alternatively, the observed changes in the amides spectral ranges could be associated with histone acetylation. Histone acetylation introduces an acetyl group (–COCH₃) on lysine residues, leading to increased intensity of CH₃ stretching and bending vibrations (∼2950–2850 cm^-^¹ and 1450 cm^-^¹respectively) and the appearance or enhancement of carbonyl (C=O) bands, which directly reflect the presence of the newly formed amide linkage. Additionally, neutralization of the lysine positive charge disrupts electrostatic interactions and hydrogen bonding within histones and between histones and DNA, resulting in shifts and intensity changes in the Amide I and Amide II bands that indicate alterations in protein secondary structure and increase of the content of unstructured turns and coils and chromatin organization [58], [59]. While blue shifts to higher wavenumber values in phosphate motions reflect conformational transitions of DNA from the B-like form to alternative structures such as A-DNA or Z-DNA, followed by chromatin damage [13], [60]. The deoxyribose-phosphate backbone can act as a hydrogen bond acceptor, and the efficiency of the hydrogen bonding formation depends on the DNA conformation. Therefore, the observed blue shifts arise from changes in hydrogen bonding between the DNA in both conformations, reflecting the reinforcement of the phosphate bonds [61]. Such molecular alterations were observed in spectra collected from chromosomes treated with 50 μM bleomycin and in cellular nuclei exposed to 150 μM bleomycin after 24 hours of treatment (Fig. 1A, C). The most intensive repair processes were detected after 48 hours of treatment in chromosomes treated with 150 μM bleomycin and in cellular nuclei exposed to 50 μM bleomycin (Fig. 1B, D). The observed differences between incubation times signify that certain repair processes in cells were partially completed between the 24 and 48-hour intervals. This observation is consistent with a report by Lipiec et al., who observed a similar effect in living cells and isolated cellular nuclei exposed to ultraviolet radiation (UVR). In SR-FTIR measurements, DNA repair processes, characterised by a partial conformational shift to A-like DNA and an increase in the amide II band, vary depending on the duration of UVR exposure [57].

The amide I region is associated with the protein backbone conformation, thereby providing insights into protein secondary structure [62]. Variations in the amide I distribution are associated with the DNA damage response, leading to increased cellular enzymatic activity and the synthesis of additional repair proteins. This chemical modification was detected in spectra from bleomycin-treated chromosomes (150 μM bleomycin dose) compared to the control group for both 24h and 48h incubation periods (Fig. 1A, 1B). Notably, there was a conformational shift from an α-helical structure in the control group to a β-sheet structure in damaged chromosomes [53]. A similar change was also observed in the spectra of cellular nuclei collected from the highest drug dose after 48 hours of treatment (Fig. 1B). Izumi et al. observed structural alterations in histone H3 proteins induced by posttranslational modifications in response to DNA damage. The authors propose that changes in the secondary structure content of H3 proteins affect the interactions of methylated histones with other molecules, such as histone-binding proteins, during DNA repair processes [63]. Therefore, we contend that the observed spectral changes in the obtained chromosome spectra reflect alterations in protein conformation occurring during the repair processes. However, to more precisely quantify these changes across the entire spectral dataset, Principal Component Analysis (PCA) was performed (Fig 3). Furthermore, local variations in the amide I to amide II ratio, which reveals changes in the relative content of secondary structures across individual chromosomes, were also examined in the later stages of this research (Fig 2).

**Figure 2.**
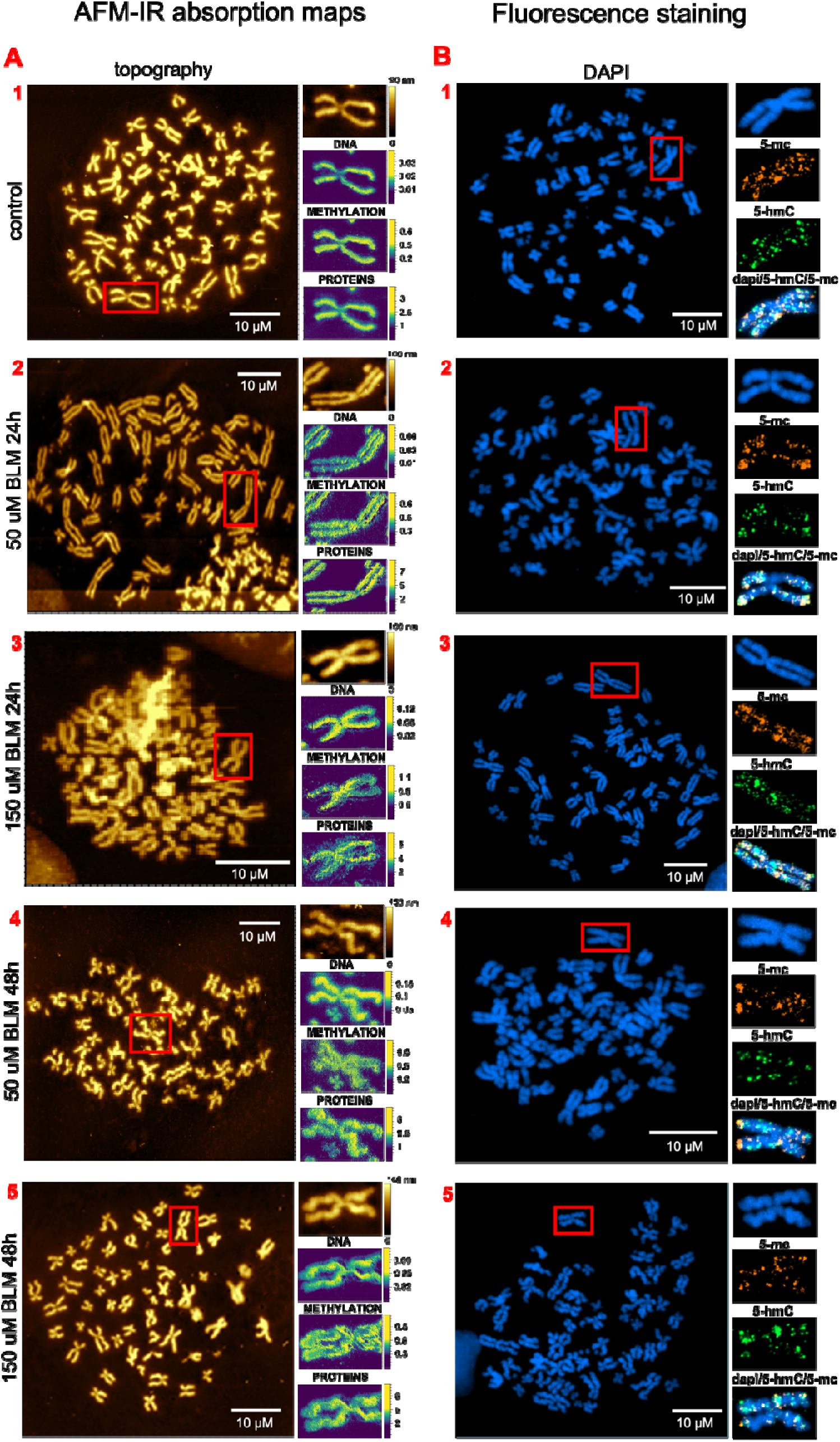
Analysis of metaphase chromosomes I identified within metaphase spreads prepared from HeLa cells treated with Bleomycin for 24 and 48 hours: (A) AFM-IR data, including: AFM topography of metaphase chromosomes with a zoomed-in area of chromosome I, presented alongside the integrated area of the absorption band at 1070 cm corresponding to the O-P-O symmetric stretching vibration; the ratio of the integrated absorption band at 1408 cm to the absorption band at 1070 cm, indicative of alterations in DNA methylation; and the ratio of the integrated absorption band at 1660 cm to the absorption band at 1540 cm reflecting changes in protein conformation. (B) HR-LSCM results, featuring metaphase chromosomes with a zoomed-in area of chromosome I, co-immunostained for DAPI (blue), 5-mc (orange) and 5-hmC (green), with images merged for comprehensive visualization.

**Figure 3.**
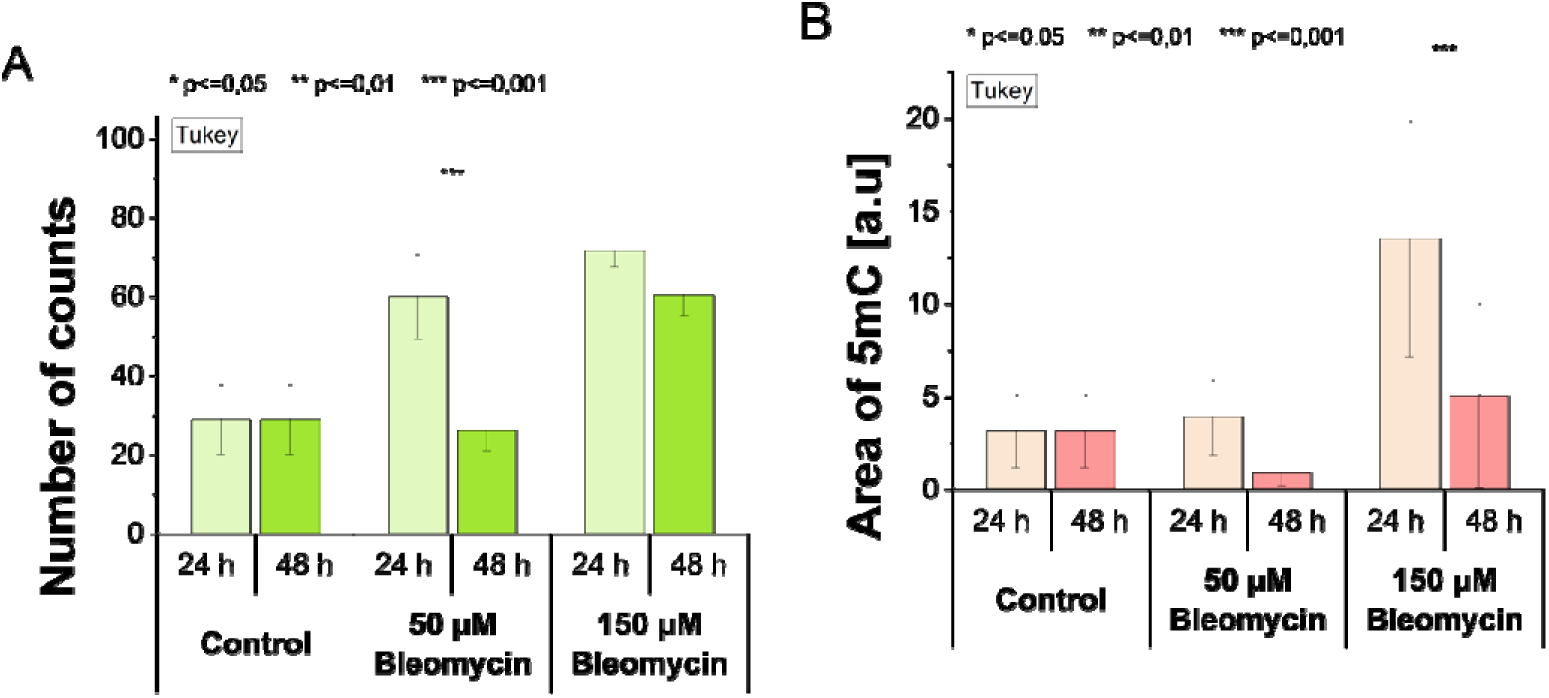
Investigation of global DNA methylation levels. (A) Semi-quantitative fluorescence intensity analysis of 5-hydroxymethylcytosine (5hmC) in metaphase chromosomes I identified within metaphase spreads prepared from HeLa cells exposed to bleomycin treatment for 24 and 48 hours at varying concentrations. (B) Comparative analysis with fluorescence staining of 5-methylcytosine (5mC).

Spectral changes observed in the region of CH_2_, CH_3_ bending and wagging mode at 1450 cm^-1^ or methyl deformation mode from cytosine at 1408 cm^-1^ can be associated with alternation in global DNA methylation and chromatin remodelling [51], [52]. In both experiments (chromosomes and cellular nuclei) for 48-hours of BLM treatment, the intensity of methyl and methylene motions decreased in relation to the applied bleomycin doses (Fig. 1B). We confirmed the decrease in global DNA methylation by 5mC and 5hmC staining (Fig. 2). In our study, chromosomes isolated from cells incubated with bleomycin for 48 hours exhibited a decrease in 5mC intensity and an increase in 5hmC intensity compared to those with shorter incubation times (24-hour treatment) and the control group (Fig. 2B). According to the studies reported by Kafer et al., high 5hmC intensity can be associated with an open chromatin conformation, while high 5mC intensity can be linked to tightly packed, repressed heterochromatin [29]. These results demonstrate that relaxation of the chromatin packaging facilitates access for repair proteins. Fortuny et al. presented evidence that pericentric heterochromatin decompacts in response to UVC damage in mouse fibroblasts. They demonstrated that chromatin decompaction is a critical step in the early stages of the DNA damage response [64].

### Nanospectroscopic mapping of metaphase chromosome methylation level and chromatin integrity in the mechanisms of DNA damage and repair

To further explore chemical modifications and chromatin integrity in the mechanisms of DNA damage and repair, the distribution of methylene, amide, and phosphate vibrations along individual metaphase chromosomes was mapped. We generated maps showing the distribution of wavenumber values representing a particular vibration in the area of interest. Each resultant map illustrates the IR absorption peak height at the specified wavenumber for each pixel. Figure 2A shows the results obtained for metaphase chromosomes I identified and analysed within metaphase spreads prepared from HeLa cells treated with a genotoxic drug with different incubation times, including AFM topographies, IR absorption maps at 1070 cm^-1^ associated with the phosphate O-P-O symmetric stretching of the DNA backbone. Methyl deformation mode from cytosine at 1408 cm^−1^ is identified as a potential marker of cytosine methylation directly related to nucleic acids [65]. To overcome the influence of chromosome thickness, the signal collected from the distribution of cytosine groups at 1408 cm^-1^ was normalised to the distribution of phosphate vibrations at 1070 cm^-1^, which is directly correlated with the density and thickness of chromatin that varies along each individual chromosome. The calculated ratios highlight regions with a higher density of methylated chromatin sites. To visualise modifications in protein conformation, the signal obtained from the distribution of amide I at 1660 cm^-1^ was divided by the distribution of amide II at 1550 cm^-1^. This approach presents local chemical insights into regions where the secondary structure of the proteins is altered, with the peak ratio also potentially influenced by the presence of extended chains [66]. Notably, post-bleomycin treatment, in comparison to untreated cells, revealed inhomogeneities in the distribution of highly methylated regions and alterations in protein secondary structure along the chromosome. Based on obtained CH3/ PO2^-^ ratios, the distribution of methylation is more homogeneous after a 50 μM dose of bleomycin (Fig. 2A.2, 2A.4) compared to a 150 μM dose (Fig. 2A.3, 2A.5), where a higher frequency of methylated areas is observed. Regarding amide I/amide II ratios, a greater degree of alterations in protein conformation is evident in chromosome structure after 48 hours of treatment (Fig. 2A.4-5), compared to untreated cells and those with a 24-hour incubation period (Fig. 2A.1-3). We assumed that observed inhomogeneities form cluster damage, indicating regions where the DNA repair process is more pronounced. These findings are further strengthened by line profile analysis extracted along the arms of selected chromosomes, presented in the Supplementary Information (Supplementary Figure S1.A). The extracted profiles reveal a direct spatial correlation between methylated regions and areas exhibiting alterations in protein conformation. Such clustered damage regions are most pronounced at 150 μM bleomycin dose (Supplementary Figure S1.A5) and are partially visible after treatment with 50 μM of bleomycin (Supplementary Figure S1.A3). In contrast, in control chromatin, DNA and protein signals closely follow chromosome topography, with no evidence of localised inhomogenities (Supplementary Figure S1.A1).

To independently verify the methylation pattern in chromosome structure, immunofluorescence staining of 5-methylcytosines (5mC), 5-hydroxymethylcytosines (5hmC), and DAPI in DNA was performed (Fig. 2B). As previously noted, bleomycin treatment of HeLa cells resulted in an increase in 5hmC and a decrease in 5mC within the chromatin structure, relative to the applied drug doses. The most significant changes were observed after 48 hours of incubation (Fig. 2B.4-5). The semi-quantitative fluorescence intensity analysis of chromosome I revealed a significant increase in 5hmC foci in response to increasing drug concentrations and incubation times, except for the 50 µM Bleomycin dose at 48 hours, where the number of 5hmC foci remained comparable to the control group (Fig. 3A). This observation is consistent with AFM-IR spectral data collected after 48 hours of drug exposure and indicate active DNA repair mechanisms (Fig. 1). In contrast, the analysis of methylcytosine staining showed no notable changes in 5mC levels as a result of Blm treatment (Fig. 3B). According to the literature, the observed accumulation of 5hmC shows active DNA demethylation, arising from the conversion of 5mC to 5hmC. Kafer et al. propose that the enhanced production of 5hmC is mediated by the TET3 enzyme during DNA repair processes [29]. It is noteworthy that each chromosome was extracted from different cells, which were at slightly different stages of metaphase. Additionally, the cells could have varying metabolic activity and chromatin content [67], [68].

### Principal Component Analysis of protein secondary structure in single metaphase chromosomes in the mechanisms of DNA damage and repair

Principal component analysis (PCA) was applied to reduce the number of collected data and explore spectral variations in protein secondary structure related to DNA damage and repair mechanisms across the entire spectral dataset. Two PCA models were calculated for AFM-IR spectra collected from single metaphase chromosomes isolated from cells incubated for 24h (Fig. 4.A, C) and 48h (Fig. 4.B, D) with different doses of genotoxic drug. Each data point corresponds to the individual spectrum acquired from a small chromatin volume localised at a single chromosome in the fingerprint region 1700-1350 cm^-1^. The 2D score plots for 24 and 48 hours, shown in Fig. 4.A and 4.B, respectively, illustrate the separation of the data into three distinct groups: the control group and two groups representing spectra acquired from chromosomes isolated from cells treated with bleomycin at doses of 50 and 150 μM. Each principal component (PC) axis visualises the distribution of the spectra. For the 24-hour treatment, the partial separation between the spectra acquired from the control group and bleomycin-treated chromosomes is visible along the PC1 component (Fig. 4.C), which explains 68% of the total variance. Some spectra acquired from bleomycin-treated chromosomes are predominantly clustered on the positive side of PC1. The loadings plot of PC1, which is positively correlated with native parallel β–sheet structure at 1626 cm^-1^ and negatively correlated with β–turn at 1662 cm^-1^ and antiparallel β–sheet structure at 1690 cm^-1^ [60]. These variations reveal changes in the protein secondary structure related to the synthesis of additional proteins with relatively high parallel β-sheet content [57]or deacetylation of histones [58]. Moreover, the PC1 loading is predominantly influenced by the band at 1530 cm^-1^ corresponding to the Amide II mode. This suggests an increase in protein concentration, a marker of elevated cellular enzymatic activity associated with DNA repair [57]. The separation along PC2 (accounting for 22% of the total variance) is not pronounced after 24h of BLM incubation.

**Figure 4.**
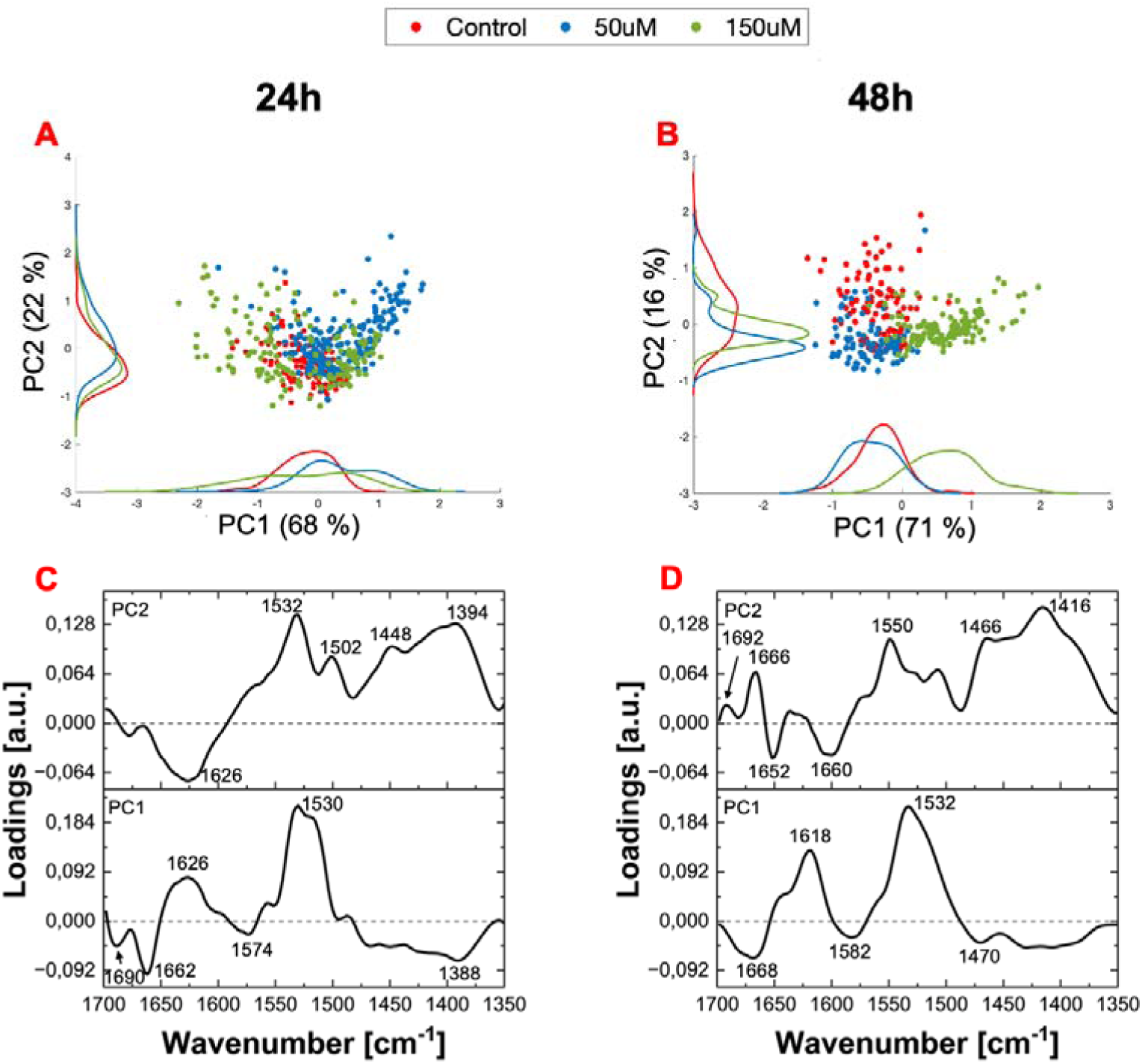
The results of PCA performed on AFM-IR spectra collected from individual metaphase chromosomes of the control group, and those incubated with 50 μM and 150 μM bleomycin for 24 and 48 hours, in the spectral range of 1700 cm to 1350 cm. (A, B) Score plots of the first two principal components (PC1 vs. PC2), each point represents an individual AFM-IR spectrum acquired along a single chromosome. (C, D) Corresponding loading plots for PC1 and PC2, indicating the spectral features that mostly contribute to the variance between the investigated groups.

The 48-hour exposure to a genotoxic drug led to a clear separation among all studied groups (control and those treated with 50, 150 µM bleomycin) in the PCA model (Fig. 4.B). Spectra acquired from 150 μM bleomycin-treated chromosomes are clustered separately on the positive side of PC1, explaining 71% of the total variance (Fig. 4.D). This loading is dominated by the maxima at 1532 cm^-1^ (Amide II vibrations) and 1618 cm^-1^ (the native parallel β-sheet structure), indicating enhanced expression of repair proteins in parallel β-like conformations that interact with DNA at the lesion site [57]. PC1 is negatively correlated with the CH2 bending mode at 1470 cm^-1^ and β-turns at 1668 cm^-1^ for the control group and 50 μM bleomycin-treated chromosomes. This result demonstrates that cells exposed to a 50 μM dose of bleomycin have largely repaired the induced DNA damage after 48h of incubation, suggesting efficient repair predominantly mediated by the NHEJ pathway [57]. The clustering of spectra, according to applied bleomycin doses, is the most evident along PC2 (representing 16% of the total variance). Most spectra obtained from chromosomes isolated from cells exposed to 50 and 150 μM bleomycin are located on the negative side of PC2, whereas over half of the spectra collected from the control group are clustered on the positive side of PC2. Such a separation is driven mainly by the PC2 loading minimum at 1652 cm^-1^ (random coil), which is a hallmark of histone acetylation, and PC2 loading maxima at 1667cm^-1^ (β–turns) and 1692 cm^-1^ (antiparallel β–sheet structure) [53], [69], [70].

Considering the results from the 24-hour and 48-hour incubation periods, the spectral changes observed reflect that for control and partially repaired cells, the amide I region is dominated by component bands at 1662-1667 cm^-1^ and 1690-1692 cm^-1^, respectively, associated with β-turns and antiparallel β-sheet structures. Conversely, for cells with active repair mechanisms, the amide I region is primarily dominated by the native parallel β-sheet structure at 1618-1626 cm^-1^.

### AFM-IR hyperspectral mapping of chromatin heterogeneity along a single metaphase chromosome

AFM-IR hyperspectral mapping was performed on a metaphase chromosome I identified within a metaphase spread prepared from HeLa cells treated with a 50 μM bleomycin dose for 24 hours, to observe local structural rearrangements in protein secondary structure and chromatin heterogeneity along the single chromosome (Fig. 5). To fully explore characteristic spectral features within the hyperspectral dataset, k-means clustering was applied to the acquired spectra. The k-means analysis identified three clusters, visualised as a false-colour map (Fig. 5B), with each colour corresponding to the average spectrum calculated for its respective cluster (Fig. 5E). The spatial distribution of identified k-means clusters clearly reflects the morphology of the mapped chromosome (Fig. 5B). However, comparison of the mean spectra revealed no significant differences in protein secondary structure. Particularly, the ratio of β-sheet structures to other secondary structure remains unchanged regardless of the distance from the centromere (Fig. 5E). Nevertheless, the relative intensity maps (Fig. B, C), highlight regions of enhanced absorption within the Amide I and Amide II regions, revealing heterogeneity in protein distribution along the chromosome. These results indicate that DNA repair processes are homogeneously pronounced in the full volume of the chromosome.

**Figure 5.**
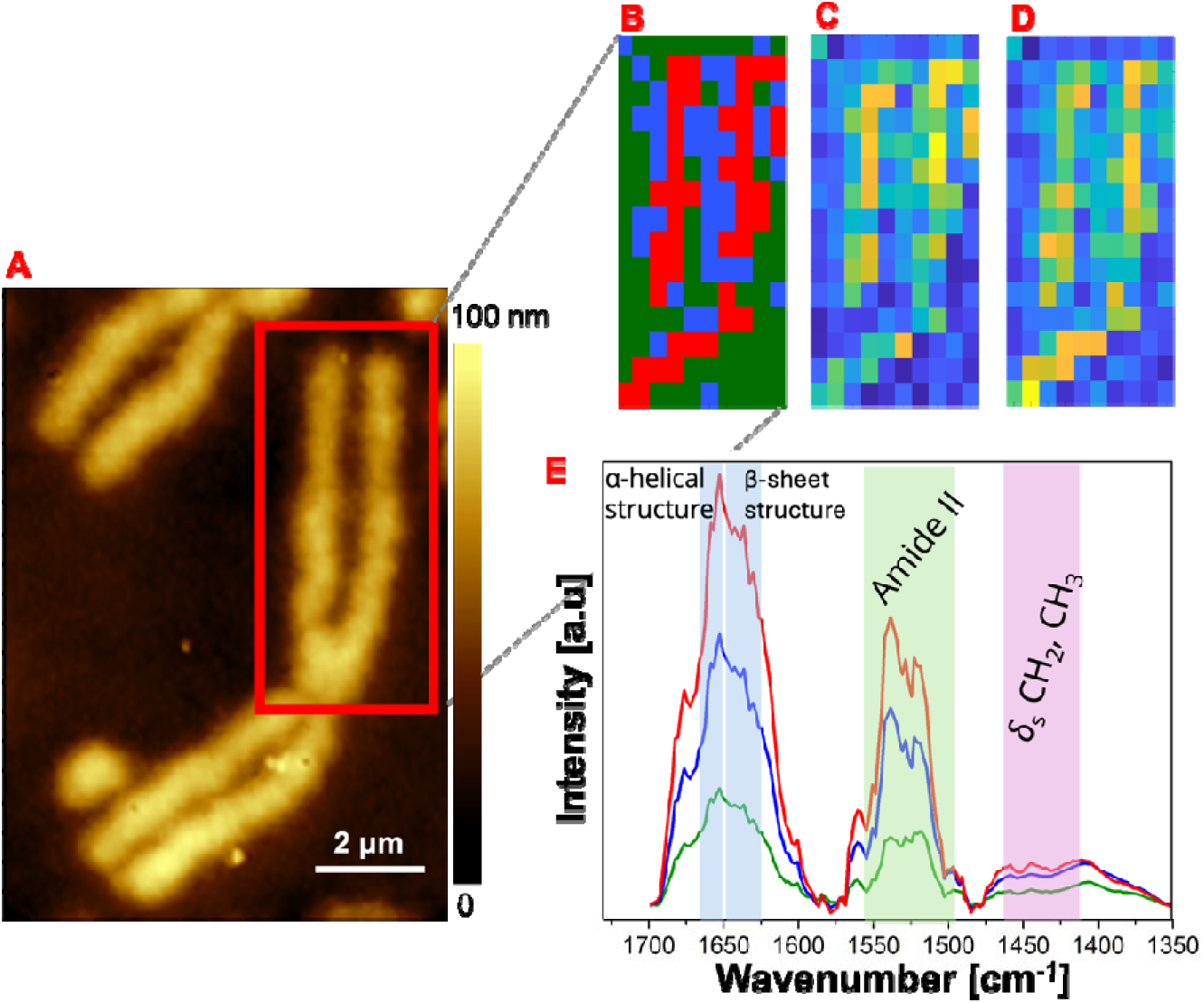
Hyperspectral mapping of a single metaphase chromosome I: (A) AFM topography of the chromosome with highlighted region indicating the area from which the map was collected. (B) The spatial distribution of the three identified k-means clusters: the green cluster corresponding to the substrate and the blue/red clusters indicating chromosome regions. (C) and (D) maps represent the relative intensity distributions for the Amide I and Amide II bands, respectively. (E) Mean spectra calculated for the three clusters identified by k-means analysis, presenting the characteristic spectral profiles for each cluster.

### Mapping of chromatin heterogeneity in chromosomal abnormalities induced by cellular responses to unrepaired double-strand breaks

We further investigated chromatin heterogeneity in chromosomal abnormalities that arise as a cellular response to unrepaired double-strand breaks, focusing specifically on the formation of dicentric chromosomes and terminal deletions, such as double minutes. Chromosomal aberrations are a form of genetic alteration resulting from incorrect repair of DSBs, leading to the fusion of two damaged chromosomes. They are associated with abnormal chromosome morphology, arising from the breakage and translocation of chromosome fragments, which are subsequently reassembled into new configurations [70], [71]. AFM topographies of investigated dicentric chromosomes and double minutes isolated from bleomycin-treated cells are shown in Figure 4A. The IR absorption maps representing the CH3/ PO2^-^ and amide I/amide II ratios show the presence of heterogeneous regions with relatively high levels of methylation and protein activity in comparison to control chromosomes isolated from untreated cells. In dicentric chromosomes, after 48 hours of treatment, a 150 μM bleomycin dose (Fig. 6.A.2) induced a higher frequency of clustered damage compared to the more homogeneous chromosome structure treated with a 50 μM bleomycin dose (Fig. 6.A.1). Accordingly, immunofluorescence staining revealed a decrease in 5mC intensity in dicentric chromosomes treated with a 150 μM bleomycin dose (Fig. 4.B.2). In the chromatin structure of double minutes, similar heterogeneities in the distribution of methylated chromatin and protein structure were observed following both 24 and 48 hours of incubation periods (Fig. 6.A.3-4). Immunofluorescence staining confirmed the presence of 5hmC foci, indicating DNA demethylation during the DNA repair process (Fig. 6.B.3-4) [29]. These observations are further supported by line profile analysis extracted along the aberrant chromosomal structures, presented in the Supplementary Information (Figure S2.A). Dicentric chromosomes present a higher frequency of cluster damage regions after 48h of incubation with 150 μM bleomycin, compared to those formed at 50 μM (Figure S2.A1-A2). In double minutes, structural inhomogeneities are visible after both 24h and 48h of treatment, indicating regions associated predominantly with NHEJ-mediated repair of DSBs (Figure S2.A3-A4).

**Figure 6.**
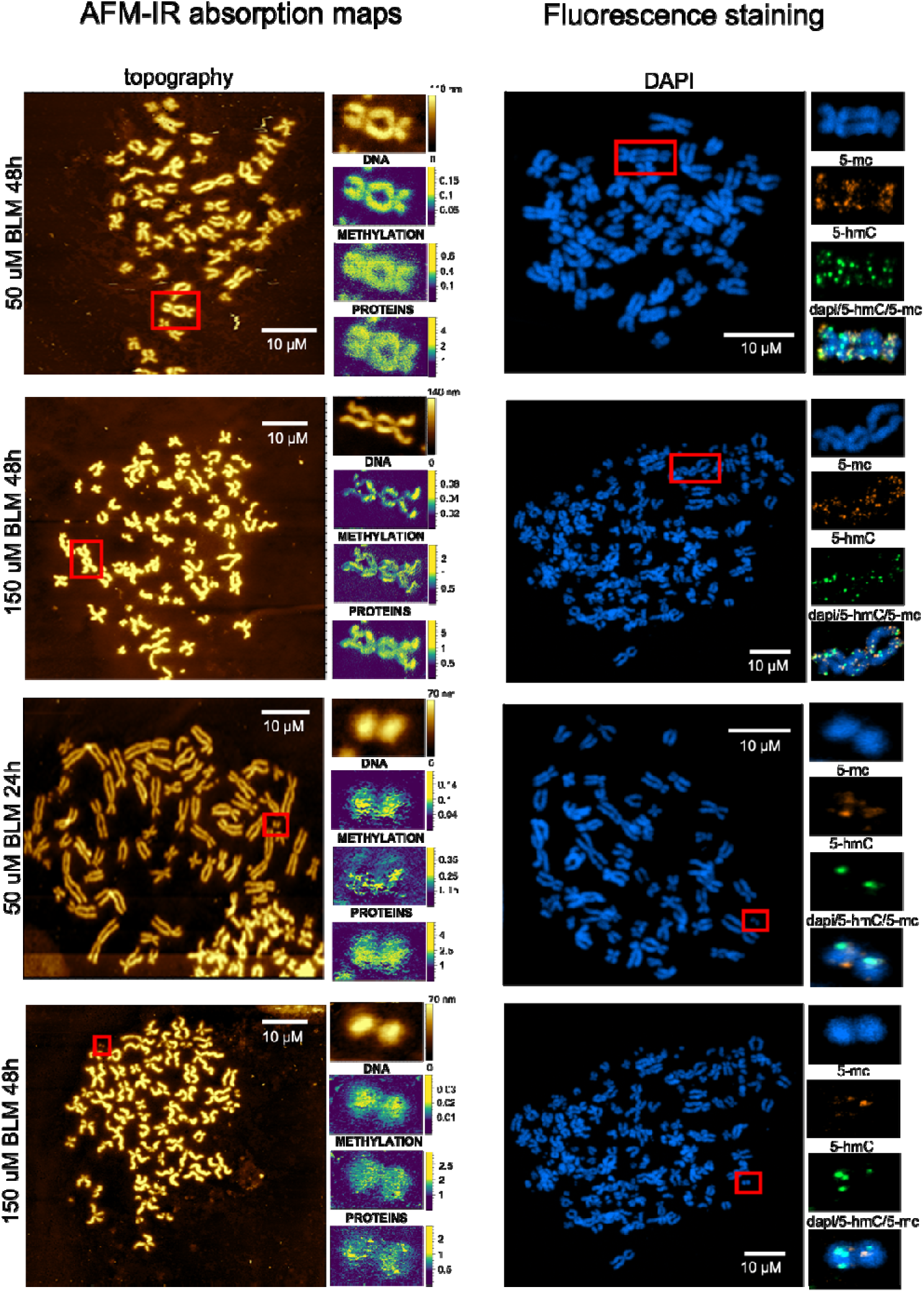
Analysis of chromosomal abnormalities induced by bleomycin treatment: (A) AFM-IR data, including: AFM topography of metaphase chromosomes with a zoomed-in area of chromosomal aberration, presented alongside the integrated area of the absorption band at 1070 cm^−1^ corresponding to the O-P-O symmetric stretching vibration; the ratio of the integrated absorption band at 1408 cm^−1^ to the absorption band at 1070 cm^−1^, indicative of alterations in DNA methylation; and the ratio of the integrated absorption band at 1660 cm^−1^ to the absorption band at 1540 cm^−1^ reflecting changes in protein conformation. (B) HR-LSCM results, featuring metaphase chromosomes with a zoomed-in area of chromosomal aberration, co-immunostained for DAPI (blue), 5-mc (orange) and 5-hmC (green), with images merged for comprehensive visualization.

### Characterisation of chromatin response to DNA double strand breaks induced by ionising radiation

To evaluate whether the nanoscale chromatin response identified in cells treated with bleomycin reflects a general mechanism of DNA damage response, we extended our study to include DSBs induced in human lymphocytes by ionising radiation (IR). Proton irradiation induces DNA damage primarily through dense ionisation tracks that generate complex and clustered DNA lesions, including DSBs [72], [73]. These lesions may arise either directly from energy deposition within the DNA molecule or indirectly through reactive oxygen species generated by water radiolysis [74].

In this study, we first examined the morphology of metaphase spreads using atomic force microscopy. Based on AFM topographies, a karyogram was constructed by distinguishing and ordering chromosomes according to the relative lengths of the p and q arms and the position of the centromere (Figure 7.A). On this basis, chromosome I was identified, and chromosomal aberrations were detected in cells exposed to a 3 Gy proton dose. To further determine the chromosomal origin of the observed aberrations, post AFM-IR measurements we performed multicolour fluorescence in situ hybridisation (mFISH) staining (Figure 7.B), which enabled precise identification of chromosomes involved in structural rearrangements and confirmation of the chromosomal composition of aberrant structures. To investigate radiation-induced chemical modifications and chromatin integrity, the distribution of methylene, amide and phosphate vibrational bands was mapped along selected chromosomes: chromosome I (labelled ‘a’ in Figure 7.A), representing a morphologically normal chromatin, and dicentric chromosome (labelled ‘f’ in Figure 7.A), representing an aberrant structure arising from misrepaired DNA damage. The IR absorption maps of the CH_3_/ PO_2_^-^ ratio and amide I intensity reveal relatively homologous distribution along the control chromosome (Figure 7.C.1). In contrast, the dicentric chromosome exhibits increased levels of methylation correlated with enhanced protein intensity, forming clustered damage regions similar to those observed in the bleomycin-DNA damage model (Figure 7.C.2). These findings are further confirmed by the line profile analysis presented in the Supplementary Information (Suplementary Figure S3). For chromosome I, the DNA and protein signals largely follow chromosome topography (Supplementary Figure S3.1), whereas in the dicentric chromosome, local inhomogenities in the form of clustered damage became apparent (Supplementary Figure S3.2). PCA was performed on AFM-IR spectra collected from chromosome I (200 spectra, labelled ‘a’ in Figure 7.A) and three dicentric chromosomes identified within the same metaphase spread (200 spectra, labelled ‘f’, ‘aa’, ‘d’ in Figure 7.A). The 2D score plot (Figure 7.D.1) shows clear separation into two groups along PC1 component (representing 72% of the total variance): spectra from chromosome I cluster on the positive side of PC1, whereas spectra from dicentric chromosomes are located on the negative side. The PC1 loading is positively correlated with bands associated with B-form DNA phosphate vibrations (∼1209 cm^-1^ and ∼1070 cm^-1^), methyl deformation vibrations (∼1437 cm^-1^), and amide II contributions (∼1516 cm^-1^), consistent with the spectral profile of structurally intact chromatin (Figure 7.D.2). In contrast, the negative PC1 loading is associated with nucleobase vibrations at 1327 cm^-1^, amide II vibrations at 1567 cm^-1^, and is dominated by amide I components at 1638 cm^-1^ and 1698 cm^-1^, attributed to disorderd/random coil and antiparallel β–sheet structures, respectively (Figure 7.D.2) [69]. Such spectral signature in the amide I may be also a hallmark of histone acetylation[58]. The separation along PC2 (accounting for 12% of the total variance) is not pronounced for the selected dataset. Collectively, these results indicate that dicentric chromosomes are characterised by a pronounced increase in disordered and antiparallel β–sheet protein conformations compared with the control chromosome I, suggesting reduced DNA repair activity and chromatin reorganisation associated with misrepaired DSBs.

**Figure 7.**
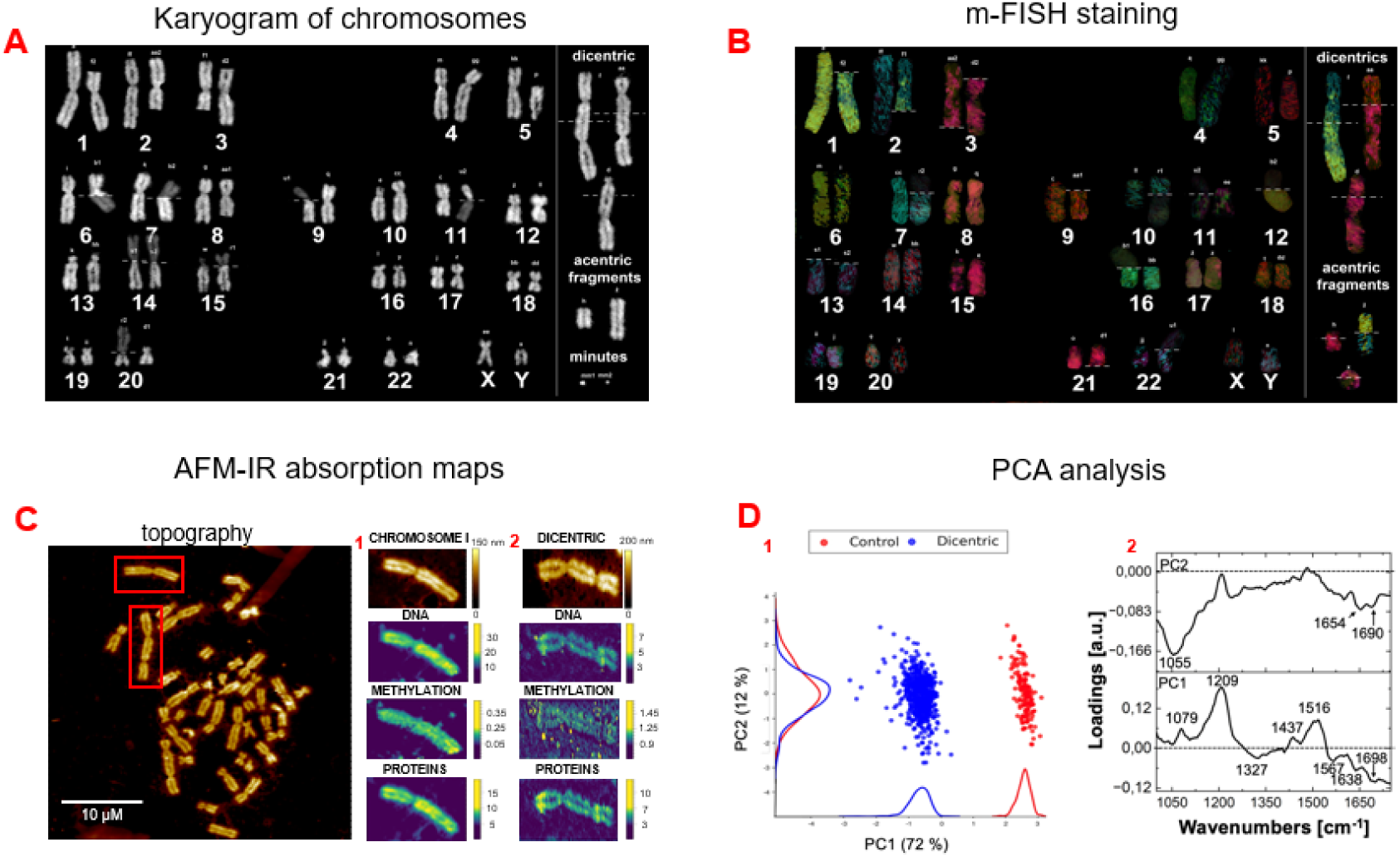
Nanospectroscopic characterisation of chromatin response to DNA damage induced by proton irradiation. (A) Karyogram of metaphase spread prepared from human lymphocytes. (B) mFISH staining confirming chromosomal identity and the structural origin of aberrations. (C) AFM topography of metaphase chromosomes with zoomed-in areas of chromosome I (C.1) and a dicentric chromosome (C.2), presented alongside the integrated area of the absorption band at 1240 cm^-1^ corresponding to O-P-O asymmetric stetching vibration of DNA backbone; the ratio of the integrated absorption band at 2960 cm^-1^ to the absorption band at 1240 cm^-1^, indicating alternations in DNA methylation; and the absorption band at 1660 cm^-1^ reflecting changes in protein secondary structure and also possibly histone acetylation. (D) The results of PCA performed on AFM-IR spectra collected from chromosome I and dicentric chromosomes within the same metaphase spread. The score plot (D.1) demonstrates separation along PC1 component, with the loading plots (D.2) highlighting contributions from DNA and protein vibrational bands driving this separation.

Considering the obtained results, the IR damage model reveals chromatin rearrangements that are consistent with those observed in the blomycin-induced DSBs model, particularly alterations in protein conformation and clustered damage accompanying misrepaired DNA breaks. These structural patterns were also investigated in oxidative DNA damage induced by hydrogen peroxide (H_2_O_2_) in chromosome sperads prepared from thyroid cancer cells, with the results presented in the Supplementary Information.

## SUMMARY AND CONCLUSIONS

In this study, we successfully combined Infrared Nanospectroscopy (AFM-IR) and high-resolution laser scanning confocal microscopy (HR-LSCM) to reveal chemical modifications in chromosome I and assess chromatin integrity following exposure to the DNA damaging agent bleomycin. The AFM-IR enabled us to monitor morphological and chemical alterations induced by the genotoxic drug, while HR-LSCM provided detailed insights into the distribution of chromatin methylation. Furthermore, we applied multivariate data analysis techniques, including Principal Component Analysis (PCA) and K-means clustering, to the AFM-IR spectra, enabling visualisation of spectral variations associated with DNA lesion induction. This multidisciplinary approach allowed us to identify specific chemical changes associated with the mechanisms of DNA damage and repair in response to bleomycin exposure.

Specifically, in chromosomes isolated from cells treated with bleomycin, we observed an enhanced expression of repair proteins accompanied by the conformational rearrangements of DNA backbone - hallmarks of double-strand break (DSB) formation and subsequent repair processes. These results are consisted with SR-FTIR spectra obtained from cellular nuclei exposed to equivalent drug concentrations and align with our previous research, where Surface-Enhanced Raman Spectroscopy revealed that DNA double-strand breaks (DSBs) induced by BLM are closely associated with conformational transitions in the DNA structure [17]. The most pronounced spectral changes were observed for chromosomes isolated from cells exposed to 50 μM bleomycin for 24 hours and 150 μM bleomycin for 48 hours. This shows that certain DNA repair processes, predominantly mediated by the NHEJ pathway, were partially completed between these time intervals. In our previous work, we have investigated individual (isolated) DNA strands. Here, we applied a more complex and biologically relevant model – integrated chromatin. Since the chromatin integrity affects the DNA ability to modify its conformation, our results from the direct nanoscale investigation of individual chromosomes upon DNA repair allowed verification whether processes more easily detectable in individual DNA strands occur also in the integrated chromatin. Therefore, our studies provide the first direct experimental verification of DNA conformational transition upon DNA repair process in individual chromosomes.

Studies of chromosomes have opened the possibility for investigating structural alterations in proteins involved in DNA repair, particularly those associated with the NHEJ pathway. Our investigations into the protein secondary structure revealed characteristic changes corresponding to DNA repair processes. PCA results indicate that in control and partially repaired cells, the amide I region is dominated by β-turns and antiparallel β-sheet structures. Conversely, in cells with active repair mechanisms, the amide I region is primarily dominated by the native parallel β-sheet structure. AFM-IR mapping further identified regions of clustered damage with increased inhomogeneities visible after 48 hours of treatment, highlighting areas where the DNA repair process is more pronounced. These findings demonstrate that protein secondary structures undergo specific local changes during the repair process.

Finally, the analysis of methylation patterns within chromosome structures showed a decrease in global DNA methylation, evidenced by the conversion of 5mC to 5hmC, a process associated with chromatin remodelling. Fluorescence staining detected the most significant methylation changes after 48 hours of drug incubation. Whereas AFM-IR CH3/ PO2-ratios indicate a higher frequency of methylated regions at a 150 μM bleomycin dose. These findings, together with previous results, reveal that chromatin relaxation enhances the accessibility of NHEJ repair proteins, thereby facilitating the DNA repair process.

Importantly, the inclusion of additional DNA damage models further strengthens the generality of our findings. Chromatin rearrangements observed in chromosomes isolated from irradiated lymphocytes revealed nanoscale alternations in protein secondary structure and local chromatin rearrangements consistent with those identified in the bleomycin model, particularly in structurally aberrant chromosomes showing signatures of reduced DNA repair activity and misrepaired DSBs. Similarly, chromosomes isolated from cells exposed to oxidative damage exhibited DNA conformational transitions, increased amide II contributions, alterations in protein secondary structure, decreased global DNA methylation, and formation of clustered damage regions.

Notably, we demonstrate that combining AFM-IR with multivariate data analysis enables the classification of chromatin from both aberrant and intact chromosomes within a single metaphase, revealing a distinct spectral signature associated with chromatin repair (Fig. 7).

We therefore conclude that the high sensitivity of our methodology opens new possibilities in chromatin research for the label-free detection of DNA damage and following the molecular modification in chromatin upon DNA repair in cases where the damage is not directly observable as chromosomal aberrations. This experimental approach has provided new molecular insights into the mechanisms underlying DNA damage and repair, highlighting the crucial roles of chromatin structure and protein conformation in these processes. These findings contribute to a deeper understanding of the cellular response to genotoxic agents and underscore the potential of nanospectroscopic techniques in the detailed investigation of chromatin structure. This comprehensive methodology can be extended in further research exploring structural and molecular alterations in chromatin integrity induced by intrinsic/extrinsic factors.

## AUTHOR CONTRIBUTIONS

Sara Seweryn, Ewelina Lipiec: conceived and planned the experiments and data analysis. Sara Seweryn, Karolina Juzba, Michał Czaja, Marta Urbańska, Anna Nowakowska: carried out the experiments. Sara Seweryn: wrote the manuscript text Katarzyna Skirlińska-Nosek: carried out the statistical analysis. All authors reviewed the obtained results and approved the final version of the manuscript.

## ACKNOWLEDGEMENTS

This work is supported by the National Science Centre, Poland under the “OPUS 16” project (Reg. No. UMO-2018/31/B/ST4/02292).

