## Supplementary Information for "Nanoscale insights into chromatin integrity molecular rearrangements upon DNA damage response"

**S1 Line profile analysis of AFM-IR absorption maps for chromosome spreads prepared from HeLa cells treated with bleomycin and human lymphocytes exposed to 3 Gy proton dose**

To complement the visual interpretation of AFM-IR absorption maps, line profiles were extracted along the arms of selected metaphase chromosomes (Figure S.1.A) and chromosomal aberrations (Figure S.2.A) prepared from HeLa cells treated with bleomycin, as well as from chromosomes isolated from irradiated human lymphocytes (Figure S3). The extracted line profiles allow direct comparison of spatial variations across different vibrational markers, revealing local associations between DNA methylation and protein conformational alterations. Signal intensities were averaged over approximately 0.4-0.8 μm along the chromosomal length. In the AFM topography images, red lines indicate the trajectories used for profile extraction, whereas red arrows in the corresponding plots highlight regions of clustered damage associated with enhanced local chromatin remodelling.

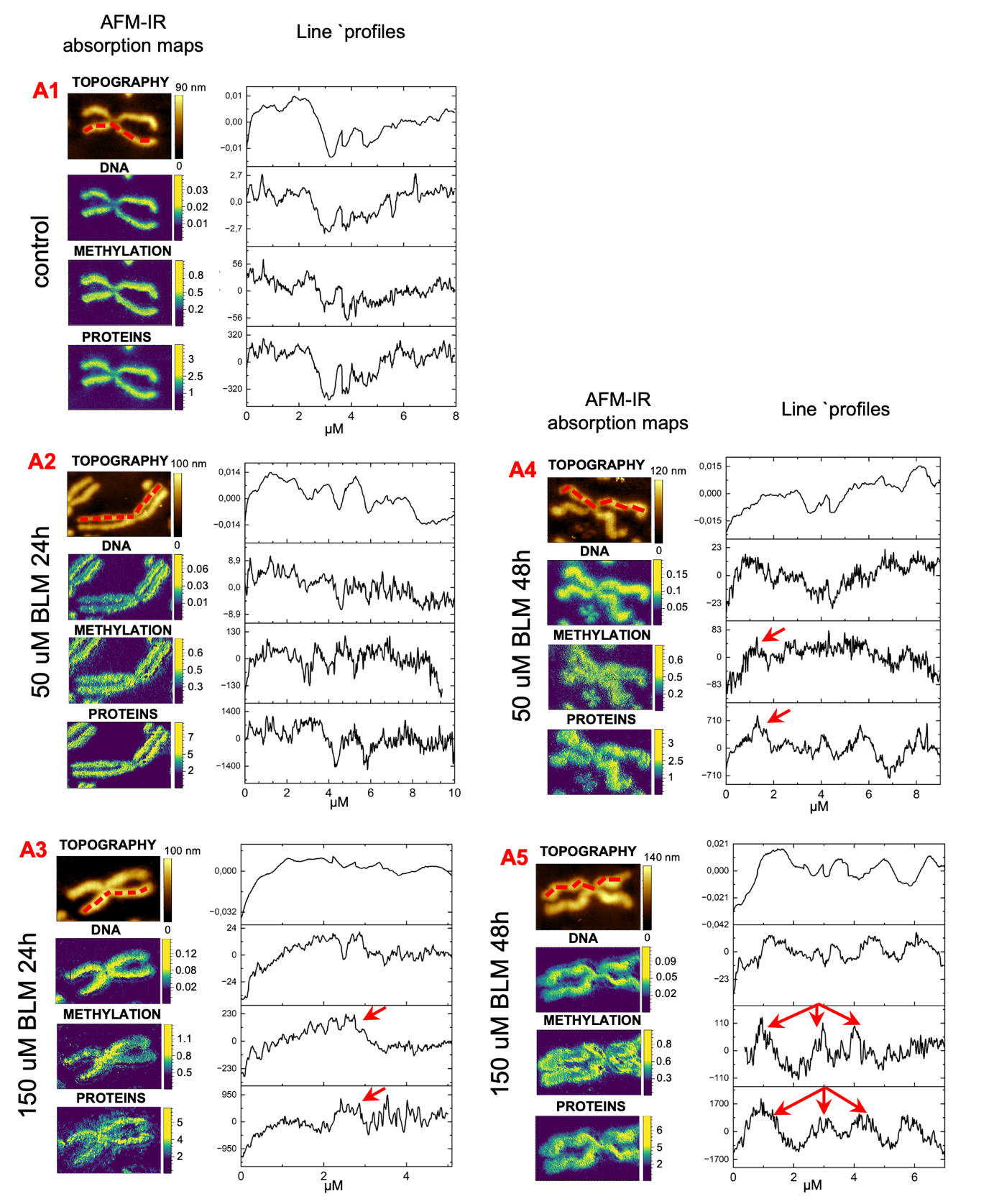

**Supplementary Figure S1.A.** Line profile analysis of AFM-IR absorption maps of chromosomes prepared from HeLa cells treated with BLM. Line profiles extracted along the arms of selected metaphase chromosomes, following the trajectories indicated by red lines in the corresponding AFM topography images. Red arrows in the profiles indicate regions of clustered damage.

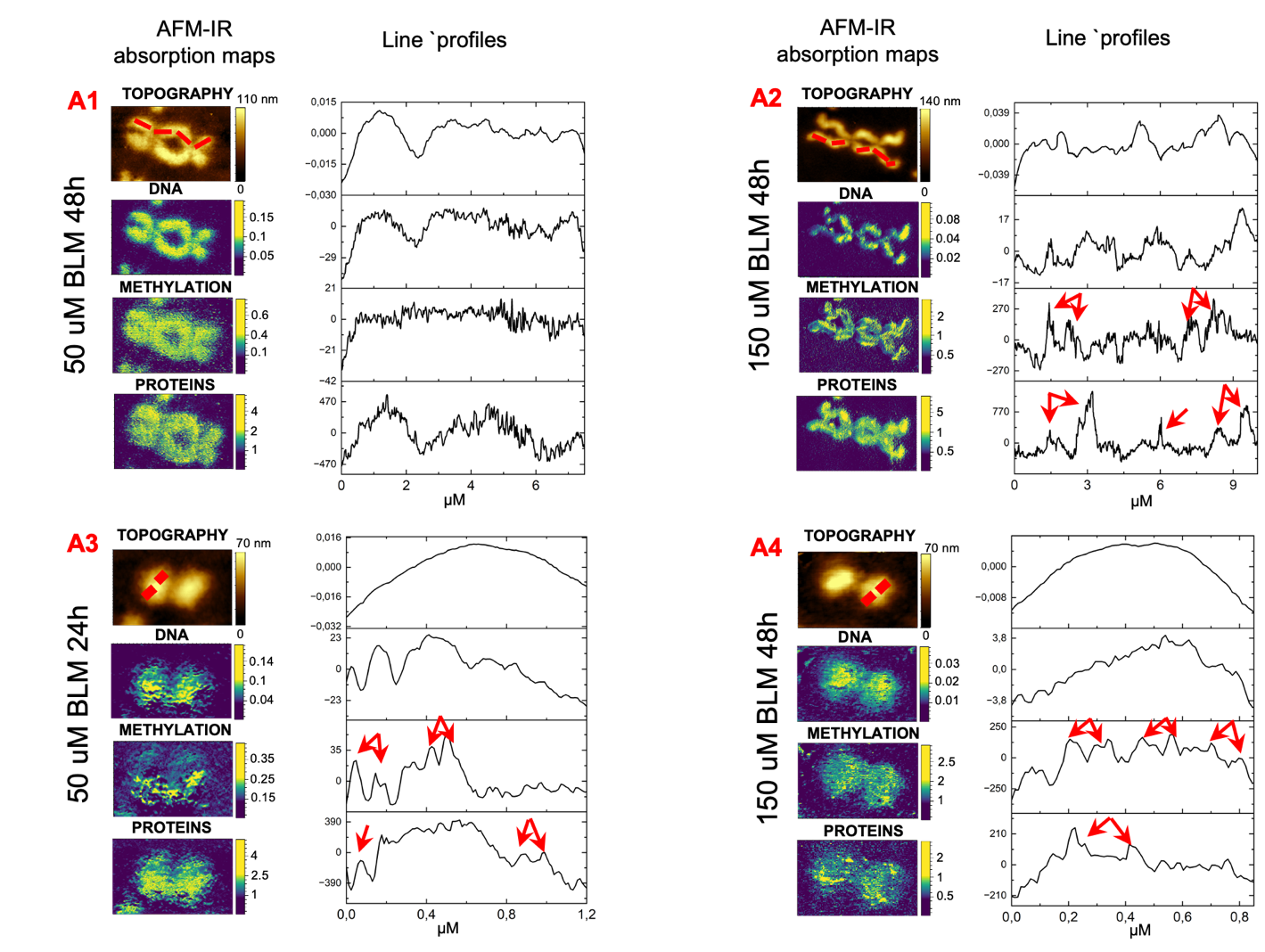

**Supplementary Figure S2.A.** Line profile analysis of AFM-IR absorption maps of chromosomal aberrations isolated from HeLa cells treated with BLM. Line profiles extracted along the arms of chromosomal aberrations following the trajectories indicated by red lines in the corresponding AFM topography images. Red arrows in the profiles indicate regions of clustered damage.

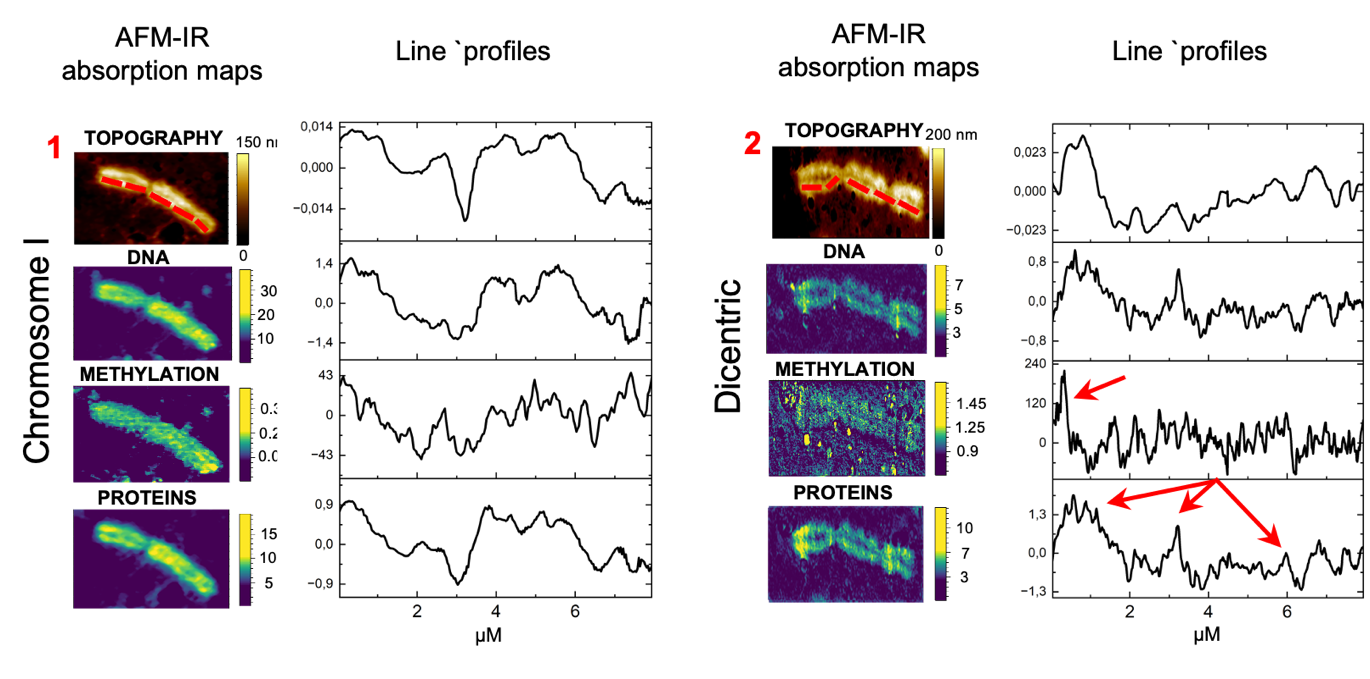
**Supplementary Figure S3.** Line profile analysis of AFM-IR absorption maps of chromosomes prepared from human lymphocytes exposed to IR. Line profiles extracted along the arms of selected chromosomes following the trajectories indicated by red lines in the corresponding AFM topography images. Red arrows in the profiles indicate regions of clustered damage.

**S2 Characterisation of chromatin response to oxidative DNA damage induced by hydrogen peroxide**

To further verify whether the nanoscale chromatin alterations identified in drug induced and radiation induced DNA damage models represent a broader cellular response, oxidative DNA damage induced by hydrogen peroxide (H_2_O_2_) in thyroid cancer cells was also investigated using the same experimental framework.

**Technical details of sample preparation and measurement procedures**

The human thyroid cancer cell line (MDA T68) was cultured at 37 °C in 5% CO₂ in high-glucose RPMI 1640 medium (Gibco) supplemented with 10% FBS (Gibco), 2 mM L-glutamine (ATCC® 30-2214™), 0.1 mM non-essential amino acids (NEAA, 100×; Gibco, cat. no. 11140050), and 1% penicillin/streptomycin (Gibco). For iron loading, a freshly prepared FeCl₂·4H₂O solution was added to the culture medium to achieve a final concentration of 0.5 µM, followed by incubation for 1 h. After PBS rinsing, oxidative stress was induced by adding hydrogen peroxide (H₂O₂) at final concentrations of 5 µM or 50 µM, followed by 48 h incubation under standard culture conditions. Metaphase chromosome spreads and isolated cellular nuclei for SR-FTIR analysis were prepared according to the protocols previously described for HeLa cells treated with bleomycin. AFM-IR and SR-FTIR measurements were performed using analogical instrumental and acquisition parameters as those used in the bleomycin experiment, as well as spectral preprocessing, infrared maps processing, PCA analysis and fluorescence staining of 5mC, 5hmC and DAPI.

**AFM-IR spectra of individual metaphase chromosomes and FTIR microspectroscopy of isolated cellular nuclei exposed to oxidative DNA damage**

AFM-IR spectra of chromosome I were compared with spectra of isolated cellular nuclei obtained using SR-FTIR at the Australian Synchrotron in Melbourne. Supplementary Figure S4 presents a comparison of AFM-IR spectra (600 averaged spectra acquired from three different chromosomes) from the control group and cells exposed to oxidative damage, with spectra of cellular nuclei (3 averaged spectra) collected using synchrotron radiation.

Hydrogen peroxide treatment induced characteristic spectral features associated with DNA repair processes. In control chromosomes, the band corresponding to antisymmetric phosphate stretching of the DNA backbone was resolved into two components, representing B-like and A-like conformations at 1201 and 1254 cm^-1^, respectively (Supplementary Figure S4.C). Following oxidative stress, a slight shift of the phosphate band toward higher wavenumbers was observed, indicating a reduced contribution of the A-like DNA conformation in treated cells and suggesting alterations in the DNA backbone. A similar, though less pronounced, effect was detected in spectra acquired from isolated cellular nuclei (Supplementary Figure S4.D). The amide II band intensity increased most prominently in chromosomes treated with 0.05 mM hydrogen peroxide, consistent with enhanced protein recruitment and active repair processes (Supplementary Figure S4.A). Comparable repair signatures were detected in SR-FTIR spectra at both applied doses (Supplementary Figure S4.B). These molecular patterns were also observed in the bleomycin model, particularly under conditions associated with active repair (Figure 1B).

Alternations in amide I distribution further revealed protein secondary structure rearrangements. A conformational shift from β-sheet toward α-helical structure was most pronounced at 0.005 mM H_2_O_2_ (Supplementary Figure S4.A). In contrast, spectra collected from cellular nuclei exhibited a partially opposite tendency (Supplementary Figure S4.B). These molecular alterations were further validated across the full dataset using PCA analyses (Supplementary Figure S8) and infrared absorption map ratio analysis (Supplementary Figure S5).

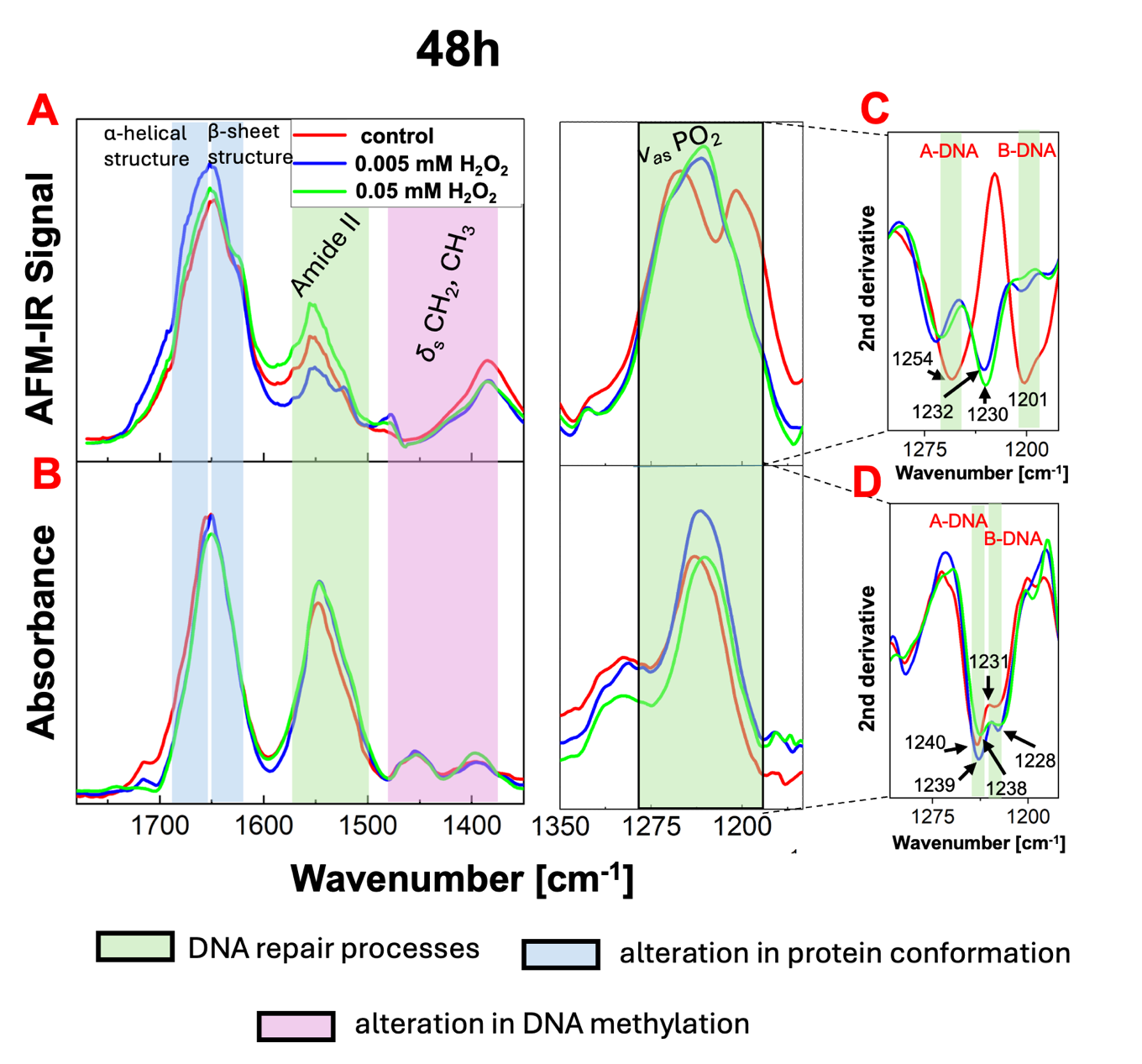
Global DNA methylation changes were also detected. In chromosome spectra, the intensity of methyl deformation vibrations at 1380 cm^-1^ decreased following hydrogen peroxide treatment (Supplementary Figure S4.A), indicating chromatin remodelling. This observation was further confirmed by 5mC and 5hmC staining (Supplementary Figure S5). A similar effect was observed in cells treated with bleomycin after 48h (Figure 1B). In cellular nuclei spectra, changes in DNA methylation were less pronounced, with a subtle increase observed at the 0.05 mM dose.

**Supplementary Figure S4.** Averaged infrared spectra illustrating DNA chemical modifications induced by hydrogen peroxide treatment. (A,B) The averaged AFM-IR spectra of single metaphase chromosomes I were compared with the averaged SR-FTIR spectra of isolated cellular nuclei after 48h H_2_O_2_ treatment. (C,D) Second derivative spectra corresponding to panels A and B, calculated to minimise baseline effects and enhance subtle spectral differences. The shaded region highlights the 1200 -1150 cm^-1^ spectral range associated with phosphate vibrations of the DNA backbone.

**Nanospectroscopic mapping of metaphase chromosome methylation level and chromatin integrity in the mechanisms of DNA damage and repair**

To assess chemical modifications and chromatin integrity, the distribution of methylene, amide and phosphate vibrations along individual metaphase chromosomes was mapped. Supplementary Figure S5.A presents the AFM topographies and IR absorption maps of chromosome I identified within metaphase spreads prepared from thyroid cancer cells exposed to oxidative stress, including the antisymmetric O-P-O stretching of the DNA backbone at 1230 cm^-1^, methyl deformation mode from cytosine at 1408 cm^-1^, and amide I, amide II bands at 1660 and 1540 cm^-1^, respectively. Ratio maps calculated using the same approach as in the bleomycin model revealed more homogenous methylation distribution in control chromosomes and those treated with 0.005 mM hydrogen peroxide (Supplementary Figure S5.A1, A2) compared with a 0.05 mM dose (Supplementary Figure S5.A3), where a higher frequency of methylated areas was observed. Protein conformational alternations, reflected by the amide I/amide II ratio, were also more pronounced following treatment with the higher H_2_O_2_ dose (Supplementary Figure S5.A3). Line profile analysis (Supplementary Figure S6) further confirmed the presence of clustered damage regions in chromosomes treated with 0.05 mM H_2_O_2_ (Supplementary Figure S6.A3), whereas in control cells and those treated with 0.005 mM hydrogen peroxide, localised inhomogeneities were less evident (Supplementary Figure S6.A1, A2). Such localised inhomogeneities were also observed in the other DNA damage models, indicating regions associated with intensified DNA repair processes.

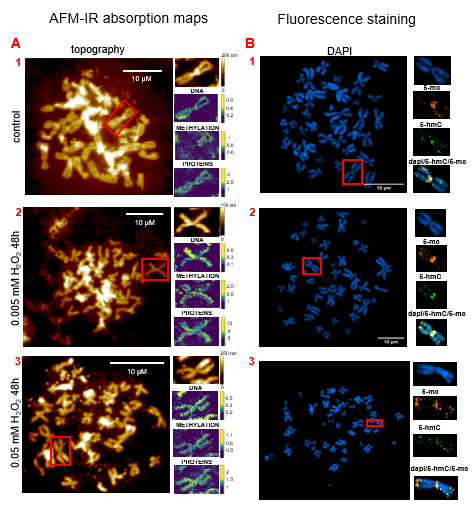

**Supplementary Figure S5.** Analysis of metaphase chromosomes I identified within metaphase spreads prepared from thyroid cancer cells treated with hydrogen peroxide for 48 hours: (A) AFM-IR data, including: AFM topography of metaphase chromosomes with a zoomed-in area of chromosome I, presented alongside the integrated area of the absorption band at 1230 cm^−1^ corresponding to the O-P-O antisymmetric stretching vibration; the ratio of the integrated absorption band at 1408 cm^−1^ to the absorption band at 1230 cm^−1^, indicative of alterations in DNA methylation; and the ratio of the integrated absorption band at 1660 cm^−1^ to the absorption band at 1540 cm^−1^ reflecting changes in protein conformation. (B) HR-LSCM results, featuring metaphase chromosomes with a zoomed-in area of chromosome I, co-immunostained for DAPI (blue), 5-mc (orange) and 5-hmC (green), with images merged for
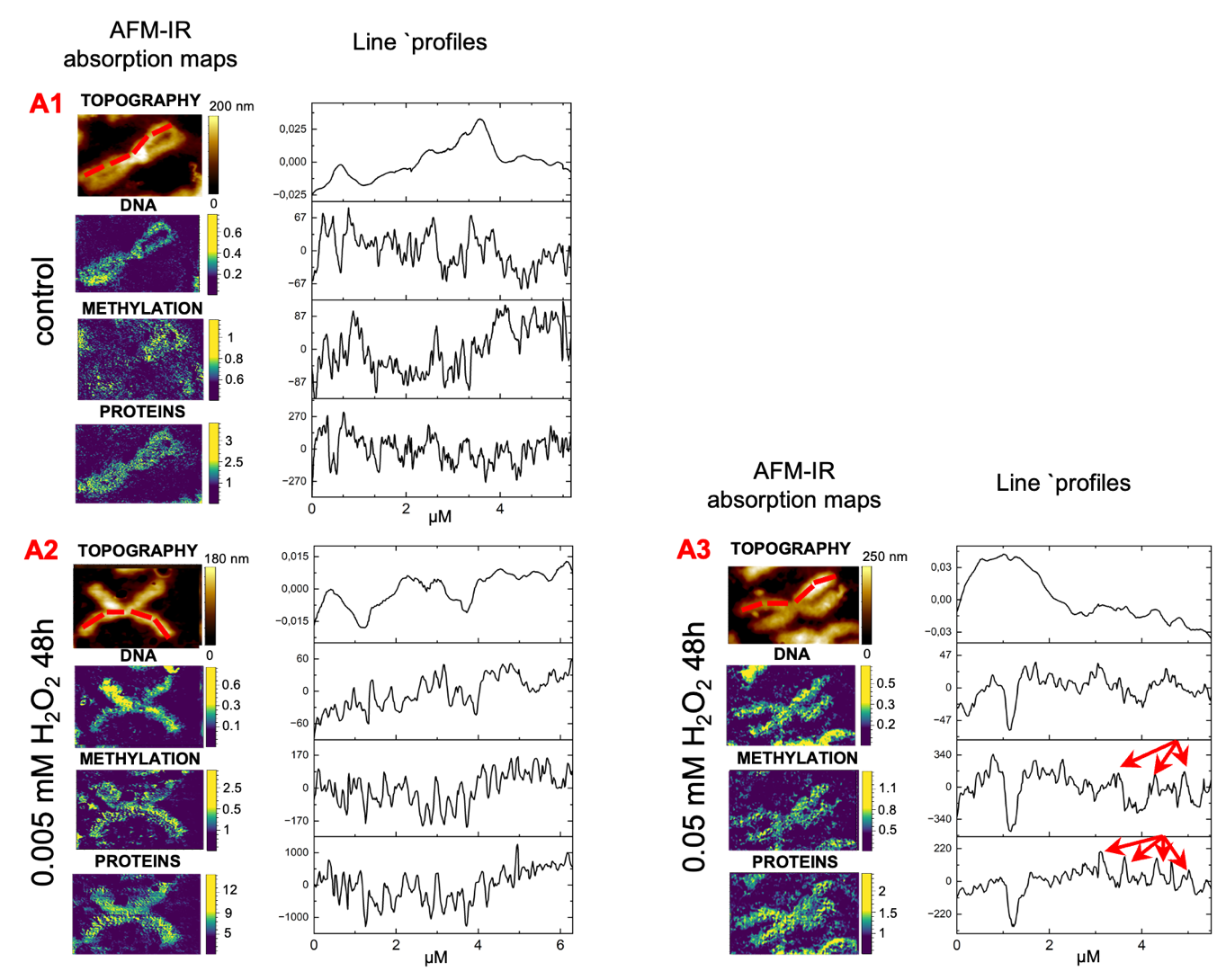
comprehensive visualisation.

**Supplementary Figure S6.** Line profile analysis of AFM-IR absorption maps of chromosomes prepared from thyroid cancer cells treated with hydrogen peroxide. Line profiles extracted along the arms of selected chromosomes following the trajectories indicated by red lines in the corresponding AFM topography images. Red arrows in the profiles indicate regions of clustered damage.

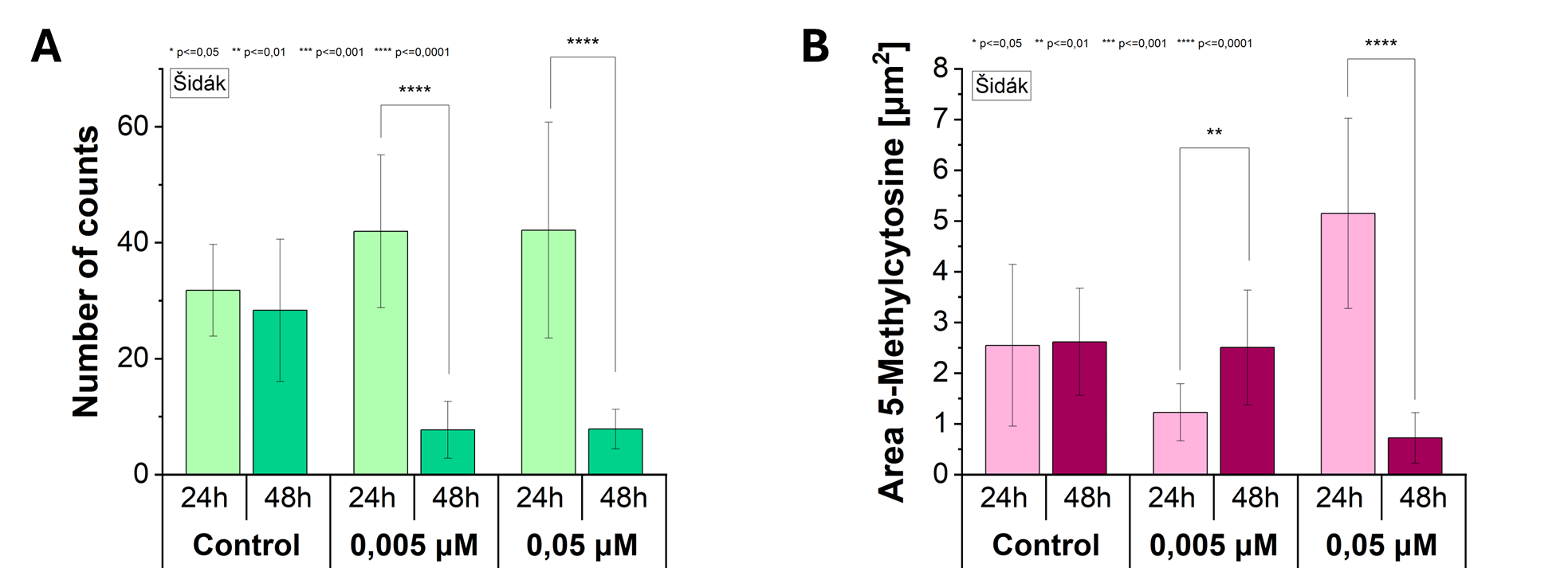

**Supplementary Figure S7.** Investigation of global DNA methylation levels. (A) Semi-quantitative fluorescence intensity analysis of 5-hydroxymethylcytosine (5hmC) in metaphase chromosomes I identified within metaphase spreads prepared from thyroid cancer cells exposed to oxidative stress for 48 hours at varying concentrations. (B) Comparative analysis with fluorescence staining of 5-methylcytosine (5mC).

**Principal Component Analysis of protein secondary structure in single metaphase chromosomes in the mechanisms of DNA damage and repair**

PCA model was calculated for AFM-IR spectra collected from single metaphase chromosomes exposed to oxidative stress for 48h. The 2D score plot shown in Supplementary Figure S8.A illustrate separation of data into two groups along the PC2 component (representing 23% of total variance). Spectra from control chromosomes and those treated with 0.05 mM H_2_0_2_ cluster on the negative side of PC2, whereas spectra from 0.005 mM dose are located on the positive side. The negative PC2 loading is associated with bands related to the DNA backbone at 1125 cm^-1^ and ~1190 cm^-1^, the Amide II mode at 1554 cm^-1^, and the native parallel β–sheet structure at 1620 cm^-1^. The positive PC2 loading is dominated by maxima at 1227 cm^-1^ (phosphate region sensitive to DNA conformation), 1270 cm^-1^ (Amide III), 1470 cm^-1^ (CH_2_ bending) and 1675 cm^-1^ (β-turns) (Supplementary Figure S8.B). These results suggest that chromosomes exposed to 0.005 mM H_2_0_2_ exhibit spectral features consistent with chromatin reorganisation under mild oxidative stress, reflected in alternations of the DNA phosphate region and increased contributions from turn-rich protein conformations. Conversely, control and 0.05 mM spectra show stronger DNA backbone and Amide II contributions, together with enhanced parallel β-sheet conformation, indicative of increased protein signal. Additional separation between control and 0.05 mM treated chromosomes is observed along the PC3 component, explaining 12% of the total variance. Spectra from chromosomes isolated from cells exposed to 0.05 mM H_2_O_2_ are located on the negative side of PC3, whereas most of the control spectra cluster on the positive side (Supplementary Figure S8.A). This separation is driven primarily by PC3 loading contributions at 1229 cm^-1^ (phosphate region sensitive to DNA conformation) and 1554 cm^-1^ (Amide II) for the 0.05 mM group, and by loading maximum at 1641 cm^-1^ (random coil) for the control group (Supplementary Figure S8.B). These findings indicate that sustained oxidative stress is associated with enhanced protein contribution and alterations in DNA conformation, whereas control chromosomes retain a more disordered protein environment.

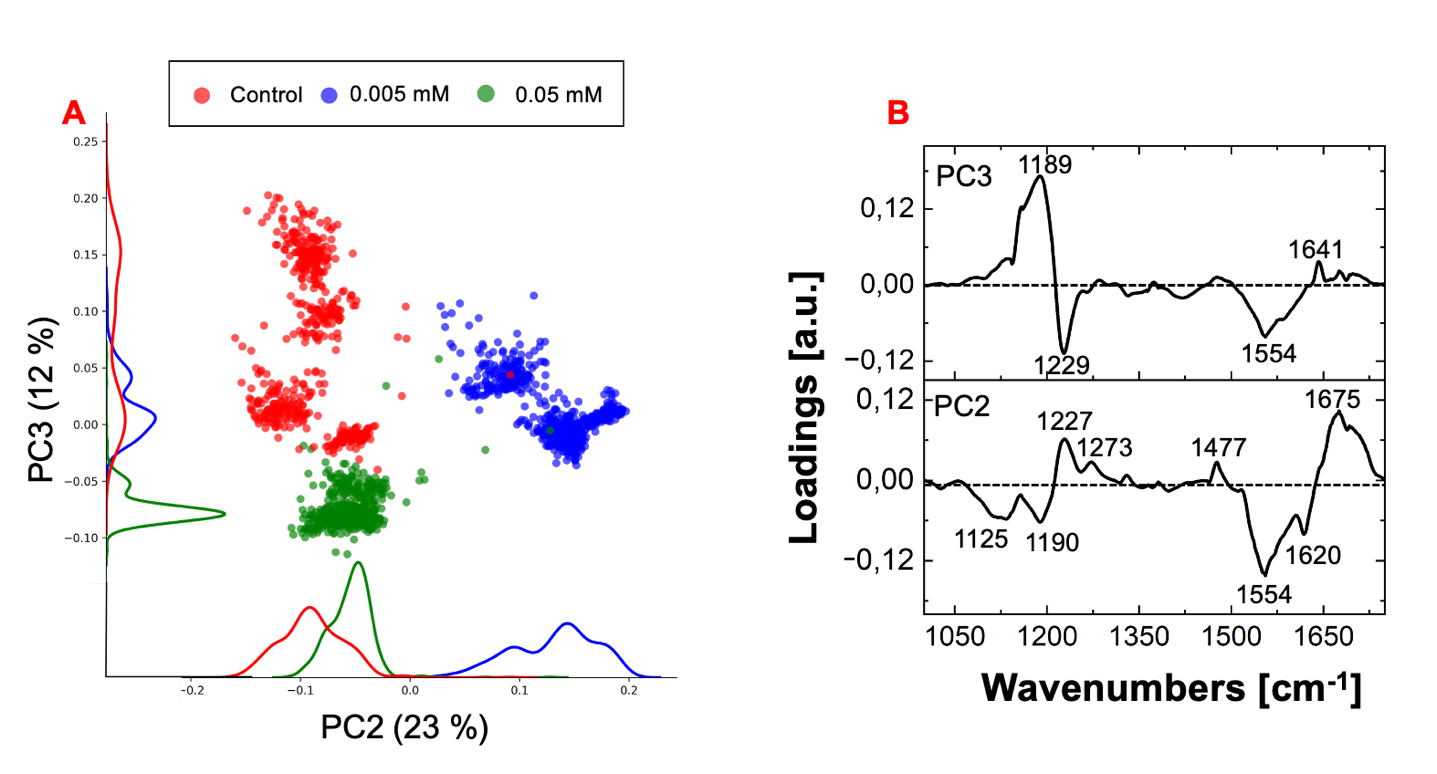

**Supplementary Figure S8.** The results of PCA performed on AFM-IR spectra collected from single metaphase chromosomes of the control group, and those incubated with 0.005 mM and 0.05 mM hydrogen peroxide for 48 hours, in the spectral range of 1700 cm^-1^ to 1000 cm^-1^. (A) Score plot of the two principal components (PC2 vs. PC3), each point represents an individual AFM-IR spectrum acquired along a single chromosome. (B) Corresponding loading plots for PC2 and PC3, indicating the spectral features that mostly contribute to the variance between the investigated groups.

**Conclusions**

The oxidative stress model confirms that the nanoscale chromatin alternations identified in the bleomycin and radiation-induced DNA damage models represent a broader and reproducible cellular response. Similar to the bleomycin model, hydrogen peroxide treatment induced characteristic spectral signatures of DNA damage and repair, including conformational transitions within the DNA backbone and increased amide II contributions indicative of enhanced protein involvement. A decrease in global DNA methylation was also observed in treated chromosomes, consistent with chromatin remodelling processes, confirmed by 5mC and 5hmC fluorescence staining. Furthermore, sustained oxidative stress led to the formation of localised clustered damage regions, suggesting areas of intensified repair activity. Alternations in protein secondary structure were also detected. Although the identified conformational patterns were different to those observed in the bleomycin and proton irradiation models. This difference likely reflects the distinct DNA repair pathway involved: bleomycin and ionising radiation predominantly induce DSBs repaired mainly through the NHEJ repair pathway[1–3], whereas oxidative DNA damage induced by hydrogen peroxide is primarily processed via base excision repair (BER), which involves a different structural organisation of repair complexes [4,5].
